# Metabolic reprogramming controlled by NF-YA alternative splicing creates therapeutic opportunities in colorectal cancer

**DOI:** 10.64898/2026.03.03.708979

**Authors:** Silvia Belluti, Valentina Mularoni, Natalia Iseppato, Virginia Campani, Mirko Ronzio, Valeria Righi, Laura Cuoghi, Arianna Rinaldi, Tommaso Martinelli, Oneda Cani, Valentina Salsi, Andrea Alessandrini, Giacomo Miserocchi, Diletta Dolfini, Vincenzo Zappavigna, Carol Imbriano

## Abstract

Metabolic reprogramming is a fundamental strategy that allows colorectal cancer (CRC) cells to endure microenvironmental constraints and sustain malignant progression. Here, we identify the transcription factor NF-Y as a master regulator of glutamine metabolism in CRC, with particular relevance to the aggressive CMS4 subtype.

Loss of function experiments, integrated with metabolomic and transcriptomic analyses, reveal a critical role for NF-YA in regulating glutamine metabolism in CRC cells. Complementary gain of function studies pinpoint NF-YAl as the isoform specifically driving glutamine-centered rewiring. Mechanistically, NF-YAl directly binds the *Glul* promoter, inducing transcriptional upregulation of glutamine synthetase and increasing intracellular glutamine availability. This metabolic reprogramming enhances resistance to mechanical shear and oxidative stress under glutamine-limiting conditions, thereby promoting migratory and metastatic traits. Importantly, pharmacological inhibition of glutamine synthesis, but not uptake or downstream catabolism, selectively abrogates the survival and migratory advantage of NF-YAl^high^ cells both *in vitro* and *in vivo*, highlighting a targetable vulnerability in aggressive CRC.

Beyond CRC cell-autonomous advantage, NF-YAl-dependent glutamine biosynthesis reshapes the tumor microenvironment by promoting M2 macrophage polarization. Conditioned medium from NF-YAl^high^ CRC cells is sufficient to induce human monocytes to adopt an M2-like phenotype. This effect is dependent on NF-YAl^high^ tumor-derived glutamine, as inhibition of glutamine uptake by monocytes fully blocks their conversion to M2. In line with this, integrative analyses of patient-derived datasets underscore the predictive relevance of the NF-YAl-*Glul*-M2 axis in driving CRC aggressiveness. These findings define glutamine synthetase as a pivotal mediator of NF-YAl activity and a promising druggable metabolic Achilles’ heel in NF-YAl^high^ CRC tumors.

## 1. Introduction

The Consensus Molecular Subtypes (CMS) and the Tumor–Node–Metastasis (TNM) systems are essential tools for risk assessment and therapeutic decision-making in colorectal carcinoma (CRC) [1,2]. Recent studies have also focused on classifying CRC patients according to their metabolic profiles [3]. Dysregulated glutamine (Gln) metabolism plays a critical role in CRC growth, metastasis, and recurrence by supporting energy production and the biosynthesis of essential macromolecules under metabolic stress [4–6]. Gln enters cells via transporters, such as SLC1A5, SLC38A1 and SLC38A2, before being converted into glutamic acid (Glu) by enzymes such as GLS1, GLS2, or GAC [7]. Glu serves multiple functions, including conversion to α-KG or participation in biosynthesis of glutathione (GSH) and nonessential amino acids (NEAAs). Mammalian cells can synthesize Gln *de novo* through glutamine synthetase (GS), encoded by the *Glul* gene. Tumor cells often overexpress GS, allowing them to reduce their dependence on external Gln sources [8–10].

We previously demonstrated that the transcription factor NF-Y regulates genes responsible for multiple metabolic pathways [11], positively controlling *de novo* biosynthetic pathways of lipids and glycolytic genes, while being neutral or repressing mitochondrial respiratory genes. NF-Y controls the SOCG (Serine, One Carbon, Glycine) and Gln pathways, as well as the biosynthesis of polyamines and purines.

Expression levels of NF-Y subunits, particularly NF-YA, are altered in multiple cancer types compared to healthy tissues [12–14]. An increase in NF-YA expression is generally associated with the up-regulation of the shorter NF-YA splice variant, NF-YAs, and a decrease in the longer isoform, NF-YAl [13,14]. Despite this, higher levels of NF-YAl characterize mesenchymal tumors and predict poor clinical outcome in breast, lung, liver, and gastric cancer patients [14–16]. As for CRC, we recently showed that increased NF-YAs is a general transcriptional hallmark of CRC compared to healthy tissues, while high NF-YAl identifies aggressive CRC characterized by EMT (epithelial to mesenchymal) and ECM (extracellular matrix) transcriptional profiles, such as MSI/CIMP and CMS4 subtypes, and predicts poor patient survival [17].

In this work, we show that NF-YA knockdown significantly alters the metabolomic profile of CRC cells. In particular, the Gln pathway is affected by NF-YA loss but enhanced by NF-YA overexpression. Gln deprivation increases NF-YAl transcription in CRC cells, while forced NF-YAl expression (NF-YAl^high^) confers resistance to Gln deprivation to cancer cells. The transcription of genes related to the Gln metabolism increases in NF-YAl^high^ cells: in particular, NF-YAl binding to the regulatory regions of *Gls1* and *Glul* genes parallels the increase in H3K4 methylation and transcription. Since blocking enzymes, transporters, or signaling pathways related to Gln metabolism represents a promising anti-cancer strategy in CRC, we explored the activity of inhibitors of the Gln pathway in NF-YAl^high^ and NF-YAs^high^ CRC cells. We demonstrate that inhibition of GS, but not of GLS or SLC1A5, reduces the aggressiveness of NF-YAl^high^ cells.

Furthermore, we show that our previously reported high metastatic potential of NF-YAl^high^ cells [17] depends, at least in part, on rewired Gln metabolism and enhanced biosynthesis of GSH, which acts as the primary intracellular antioxidant [18–21]. Such metabolic reprogramming also shapes the tumor microenvironment (TME) by influencing immune cell behavior. In particular, increased Gln consumption by tumor-associated macrophages (TAMs) drives M2 polarization, promoting immune evasion and metastasis in CRC [22]. Consistently, we found that high NF-YAl levels correlate with M2 macrophage infiltration in tumors and can directly induce M2 macrophage activation *in vitro*. Taken together, our findings suggest that inhibiting Gln synthesis in aggressive NF-YAl^high^ tumors could be a promising therapeutic strategy to limit CRC progression and improve patient outcomes.

## 2. Results

### 2.1 NF-YA depletion transcriptionally remodels metabolic programs and reshapes the CRC metabolome

To assess whether NF-Y influences CRC cell metabolism, we silenced NF-YA in HCT116 cells *via* shRNA lentiviral transduction, which severely impaired cell viability (Fig. 1A), and performed comprehensive metabolomic profiling using 1D and 2D HR-MAS NMR (Fig. 1B). Representative spectra and identified metabolites are shown in Suppl. Fig. 1 and Suppl. Table 1. Quantitative comparison between NF-YA^KD^ and control (CTR) cells revealed marked remodeling of the metabolome following NF-YA depletion (Fig. 1B). In particular, NF-Y loss significantly reduced Glutamate, N-Acetylaspartate, Pyruvate, Aspartate, Glutathione, Glycine, Threonine, Phosphatidylcholine, Taurine and Creatine levels, while Lactate displayed a non-significant upward trend, suggesting a potential shift in metabolic flux.

**Figure 1.**
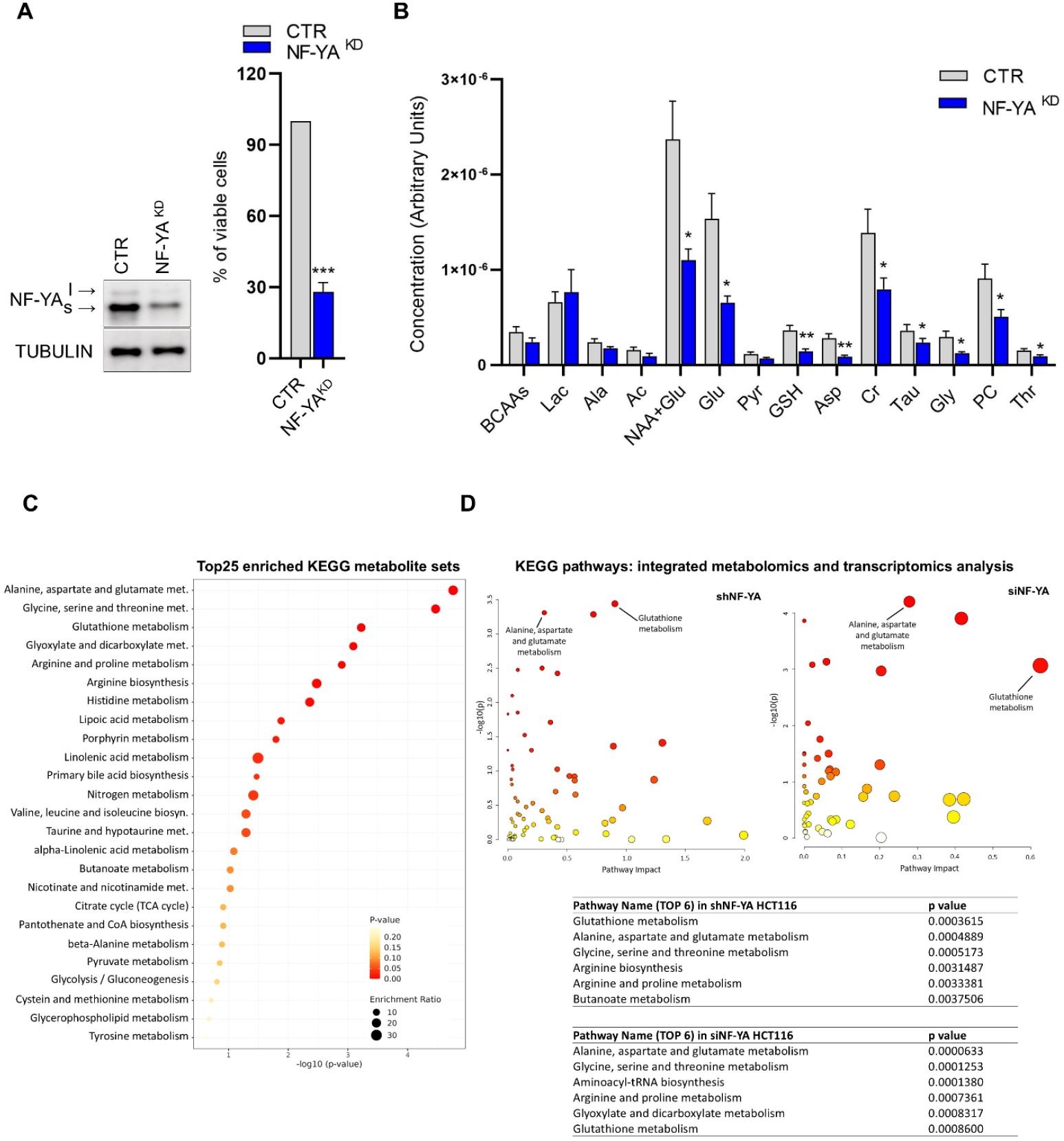
NF-YA knockdown affects cell metabolism in CRC cells. **A)** Left: Western blot analysis of NF-YA expression in whole cell extracts of shControl (CTR) and NF-YA-inactivated (NF-YA^KD^) HCT116 cells. Tubulin was used as loading control. Right: Effect of NF-YA knock-down on HCT116 cell viability compared to shControl (CTR) cells, arbitrarily set at 100%. Data represent mean±SEM (one-sample t-test, n=4) **B)** Metabolic profile obtained by HR-MAS NMR in HCT116 cells 48h-post infection with shControl (CTR) or shNF-YA (NF-YA^KD^) lentiviral vectors. Data were analysed by spectral deconvolution, normalization and fold-change analysis of the area under the receiver-operating curve. Relative concentration (Arbitrary Units) was defined as the ratio of the relative resonance area of all the quantifiable metabolites and the respective cells number. Data represent mean±SEM (t-test, n=5). **C) and D)** Results of pathways analysis based on KEGG human metabolic pathways performed with MetaboAnalyst 6.0. C) Dot plot of metabolite set enrichment analysis (MSEA) for metabolites significantly different between CTR and NF-YA^KD^cells. The size of the circles indicates the Enrichment Ratio, computed by observed hits / expected hits, while the color and x-axis represent the p-value. D) Integration of metabolites significantly different between CTR and NF-YA^KD^ cells and datasets of differentially expressed genes in CTR vs NF-YA^KD^ cells obtained from our shRNA-mediated (GSE70543) or siRNA-mediated (GSE56788) NF-YA^KD^ in HCT116 cells. The scatter plots represent matched pathways according to the p values from the pathway enrichment analysis and pathway impact values from the pathway topology analysis. The color and size of the circles indicate their p-value and pathway impact value, respectively. Gln-related pathways are indicated in the plots. The top 6 matched pathways and their p-values were also reported in the tables.

To identify metabolic pathways regulated by NF-Y, we performed metabolite set enrichment analysis (Fig. 1C) followed by integrated joint pathway analysis combining metabolomics with our matched transcriptomic dataset from shRNA-mediated NF-YA^KD^ HCT116 cells (GSE70543) (Fig. 1D). Both analyses highlighted “Alanine/aspartate/glutamate”, “Glutathione metabolism” and “Glycine/serine/threonine metabolism” as the most significantly enriched KEGG pathways. These findings were independently validated using a second transcriptomic dataset of siRNA-mediated NF-YA^KD^ cells (GSE56788) (Fig. 1D).

These findings underscore the critical role of NF-Y in metabolic homeostasis of CRC cells, with glutamine metabolism and related pathways, including glutathione and glutamate metabolism, being highly dependent on NF-YA transcriptional activity. Collectively, our results highlight a tight link between NF-YA expression and the metabolic programs that sustain CRC cell survival and proliferation.

### 2.2 NF-YA splicing modulates functional live cell metabolism and transcriptionally reshapes glutamine utilization

Transcriptional regulation of metabolic pathways profoundly affects cell growth and function. NF-Y has emerged as a key metabolic regulator [23], and we showed that the NF-YA splicing signature correlates with CRC aggressiveness, with high NF-YAl levels associated with shorter patient survival [17]. To dissect the roles of NF-YA isoforms in metabolism, we used previously characterized HCT116 cells, namely NF-YAl^high^ and NF-YAs^high^, which express high levels of NF-YAl and NF-YAs isoforms, respectively [17] (Fig. 2A). Live-cell Seahorse metabolic assays were employed to assess oxidation of glucose (Gluc), Glutamine (Gln), and long-chain fatty acids (FA). Sequential inhibition of metabolic pathways allowed evaluation of mitochondrial dependency, capacity, and flexibility for each fuel (Fig. 2B, Suppl. Fig. 2), revealing the impact of NF-YA splice variants on cellular energy metabolism. NF-YAl^high^ cells significantly increased Gln capacity and dependency compared to control and NF-YAs^high^ cells (Fig. 2B). The higher Gln capacity in NF-YAl^high^ *versus* control cells (31.8% *vs* 13.9%) indicates that Gln supports mitochondrial energy production when Gluc and FA pathways are blocked. Likewise, NF-YAl^high^ cells rely more on Gln oxidation to sustain basal respiration (19.9% *vs* 8.9% in control cells). Nevertheless, their mitochondria can compensate for Gln inhibition by using Gluc and FA, whereas NF-YAs^high^ cells exhibit markedly reduced flexibility (3.7% *vs* 12.0% in controls) (Fig. 2B).

**Figure 2.**
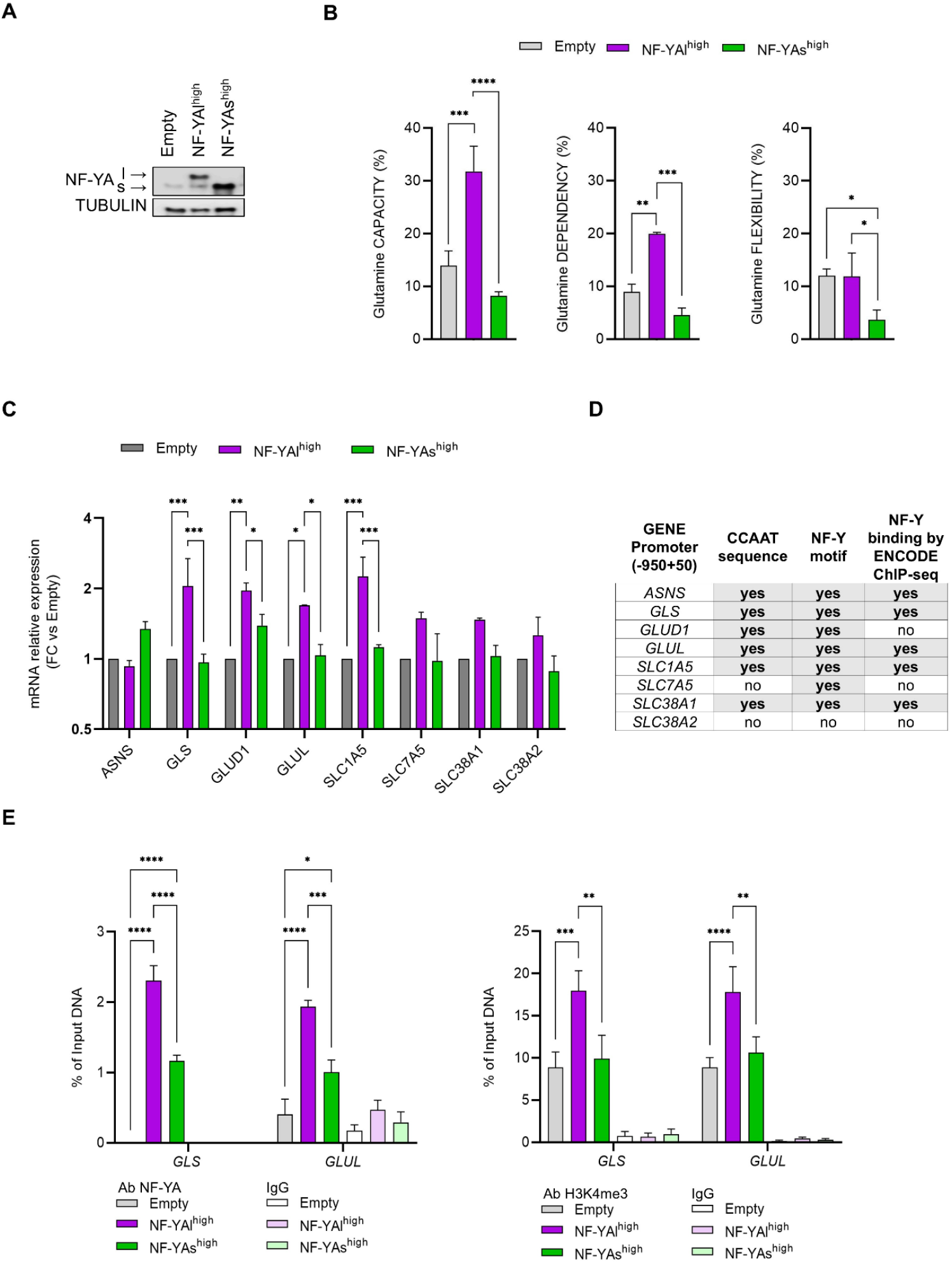
Higher expression of NF-YAl alters the metabolism of CRC cells and the expression of genes related to glutamine metabolism. **A)** Western blot analysis of total extracts from HCT116 cells stably infected with Empty, NF-YAl and NF-YAs lentiviral particles. Tubulin was used as loading control. **B)** Effect of overexpression of NF-YA isoforms on Gln capacity, dependency and flexibility (%). Data are expressed as mean±SEM (two-way ANOVA with Fisher’s LSD test, n = 3). **C)** RT-qPCR analysis of genes related to Gln-metabolism in Empty, NF-YAl^high^, and NF-YAs^high^ HCT116 cells. Rps20 was used as reference gene and normalized mRNA levels are reported as fold change vs Empty, arbitrarily set at 1. Data represent mean±SEM (two-way ANOVA with Fisher’s LSD test, n = 3). **D)** In silico analysis of selected promoters (−950bp to +50bp relative to TSS) for the presence of the CCAAT pentanucleotide or putative NF–Y transcription factor binding sites (Jaspar and Transfac matrices). The right column indicates the presence (yes or no) of NF-Y binding in ChIP-seq dataset from the ENCODE project (UCSC Genome Browser, human GRCh38/hg38). **E)** qChIP analysis of NF-YA binding (left panel) and H3K4me3 open chromatin mark (right panel) at promoter regions of *Gls1* and *Glul* genes in HCT116 cells. IgGs were used as negative control antibody. Data represent mean±SEM (two-way ANOVA with Fisher’s LSD test; 3≤n≤7).

To better understand how NF-YAl^high^ cells modulate Gln production and utilization, we analyzed key genes controlling the Gln pathway by RT-qPCRs. NF-YAl^high^ cells showed a significant increase in the transcription of Gln transporters (*Slc1a5, Slc7a5* and *Slc38a1* genes) and of key enzymes related to the Gln pathway (*Glul, Glud1* and *Gls* genes) (Fig. 2C). This finding aligns with previous transcriptomic analyses conducted on NF-YA silenced cells (Fig.1D), where Gln transporters, *Glul* and *Gls* were identified as genes affected by NF-YA knock down (Suppl. Table 2). To understand whether NF-Y directly controls the expression of Gln-related genes, we initially explored the presence of NF-Y binding sites (CCAAT/ATTGG or Transfac/Jaspar matrices) and of NF-Y binding within their regulatory regions (−450/+50 bp from the TSS) by using LASAGNA-Search 2.0 [24] and consulting ENCODE data (Fig. 2D). We then performed chromatin immunoprecipitations with anti-NF-YA and anti-H3K4me3 histone mark antibodies to identify NF-Y binding to putative NF-Y target genes in HCT116 cells (Fig. 2E). Remarkably, we identified NF-YA binding on the regulatory regions of *Glul* and *Gls*. The augmented binding of NF-YA, coupled to increased trend of positivity of the H3K4 histone mark, supported the enhanced transcription of these genes in NF-YAl^high^ cells.

Overall, our data indicate that increased basal expression of Gln-metabolism genes, directly regulated by NF-Y and supported by higher NF-YAl levels, may enhance cancer cells’ adaptability to fluctuations in Gln availability.

### 2.3 Glutamine deprivation modulates the expression of NF-YA isoforms

In CRC, low systemic Gln levels have been associated with increased tumor aggressiveness [25]. We therefore investigated the role of NF-YA isoforms in metabolic adaptation and tumor progression. To assess whether Gln availability influences the expression of NF-YA isoforms, HCT116 and SW480 CRC cells were cultured in Gln-depleted medium for 72 hours.

Under control conditions, both cell lines exhibited high NF-YAs and low NF-YAl expression (Fig. 3A, Suppl. Fig. 3A). Following Gln deprivation, mRNA and protein levels of both isoforms increased (Fig. 3A, B and Suppl. Fig. 3A, B), although the NF-YAs/NF-YAl ratio decreased markedly, from 7.80 to 2.55 in HCT116 cells (Fig. 3A) and from 8.46 to 2.33 in SW480 cells (Suppl. Fig. 3A). This shift was accompanied by a significant reduction in cell proliferation over 72 hours, which persisted up to 7 days (Fig. 3C, Suppl. Fig. 3C).

**Figure 3.**
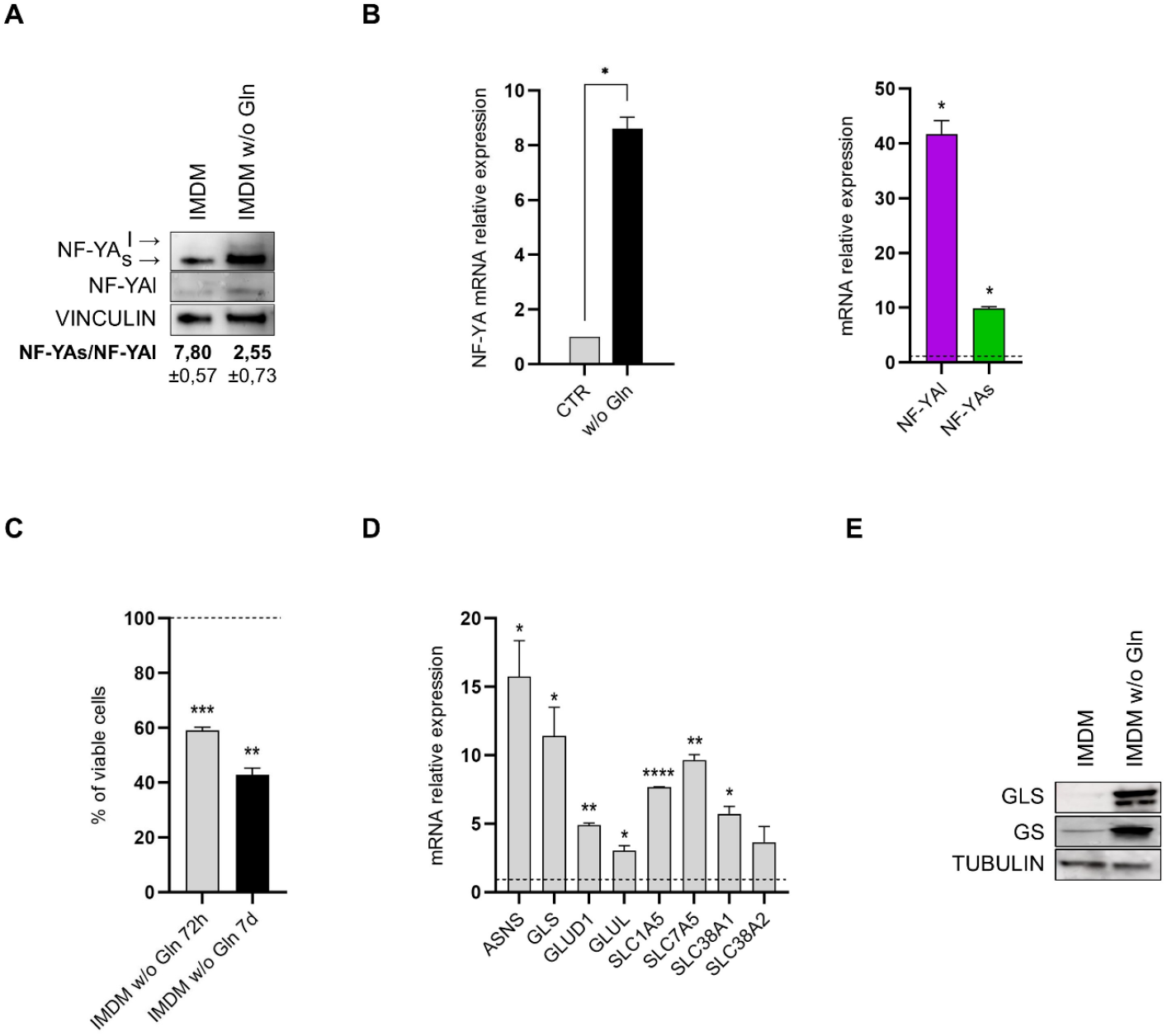
Glutamine deprivation alters the expression of NF-YA isoforms and genes related to glutamine metabolism. **A)** Protein levels of NF-YA measured by Western blot in HCT116 cells in complete IMDM medium or IMDM without Gln. Vinculin was used as loading control. The quantification of NF-YAs and NF-YAl expression by ImageJ analysis and the NF–YAs/NF–YAl ratio is reported. **B)** Transcript levels of NF-YA (left) and NF-YAl and NF-YAs splice variants (right) in complete IMDM (CTR) or IMDM w/o Gln, quantified by RT-qPCR. Results are reported as fold change vs CTR, arbitrarily set at 1 (dotted line in right panel). Rps20 was used as reference gene. Data represent mean±SEM (one-sample t-test, n=3). **C)** Cell viability of HCT116 cells analyzed by PrestoBlue cell viability assay after 72h and 7 days in Gln-deprived medium. Data were normalized to cell growth in CTR medium, arbitrarily set at 100% (dotted line), and represent mean±SEM (one-sample t-test, n=3). **D)** Transcript levels of genes involved in Gln metabolism, quantified by RT-qPCR. Results are reported as fold change vs CTR medium, arbitrarily set at 1 (dotted line). Rps20 was used as reference gene. Data represent mean±SEM (one-sample t-test, n=3). E) Protein levels of GLS and GS measured by Western blot in HCT116 cells in complete IMDM medium or IMDM without Gln. Tubulin was used as loading control.

In parallel, Gln deprivation significantly upregulated the transcriptional levels of membrane transporters and metabolic enzymes mediating both Gln catabolism and anabolism, including *Gls* and *Glul*, respectively (Fig. 3D). Consistently, protein levels of *Glul*-encoded GS and *Gls*-encoded Glutaminase also increased. These findings suggest that higher NF-YAl levels may contribute, through direct and indirect mechanisms, to the cellular response to Gln deprivation, likely by promoting the expression of genes regulating this essential metabolic pathway.

### 2.4 The inhibition of the glutamine synthetase enzyme reduces NF-YAl-associated aggressiveness of CRC cells

Since targeting Gln metabolism has emerged as a therapeutic strategy for cancer [26], and NF-YAl^high^ cells appear to be in a pre-adapted state that may confer resistance to variations in Gln availability, we evaluated the effect of Gln deprivation for 72 hours on cell growth of isogenic HCT116. Compared to control cells in Gln-supplemented medium, the viability of Empty and NF-YAs^high^ cells significantly decreased by about 34% and 45%, respectively, while NF-YAl^high^ cells showed higher resistance, with only a 10% reduction in viability (Fig. 4A). Similar results were obtained in SW480 cells (Suppl. Fig. 3D). As a control, we added back Gln to the culture medium and this either fully or nearly completely restored cell viability, as expected (Fig.4A).

**Figure 4.**
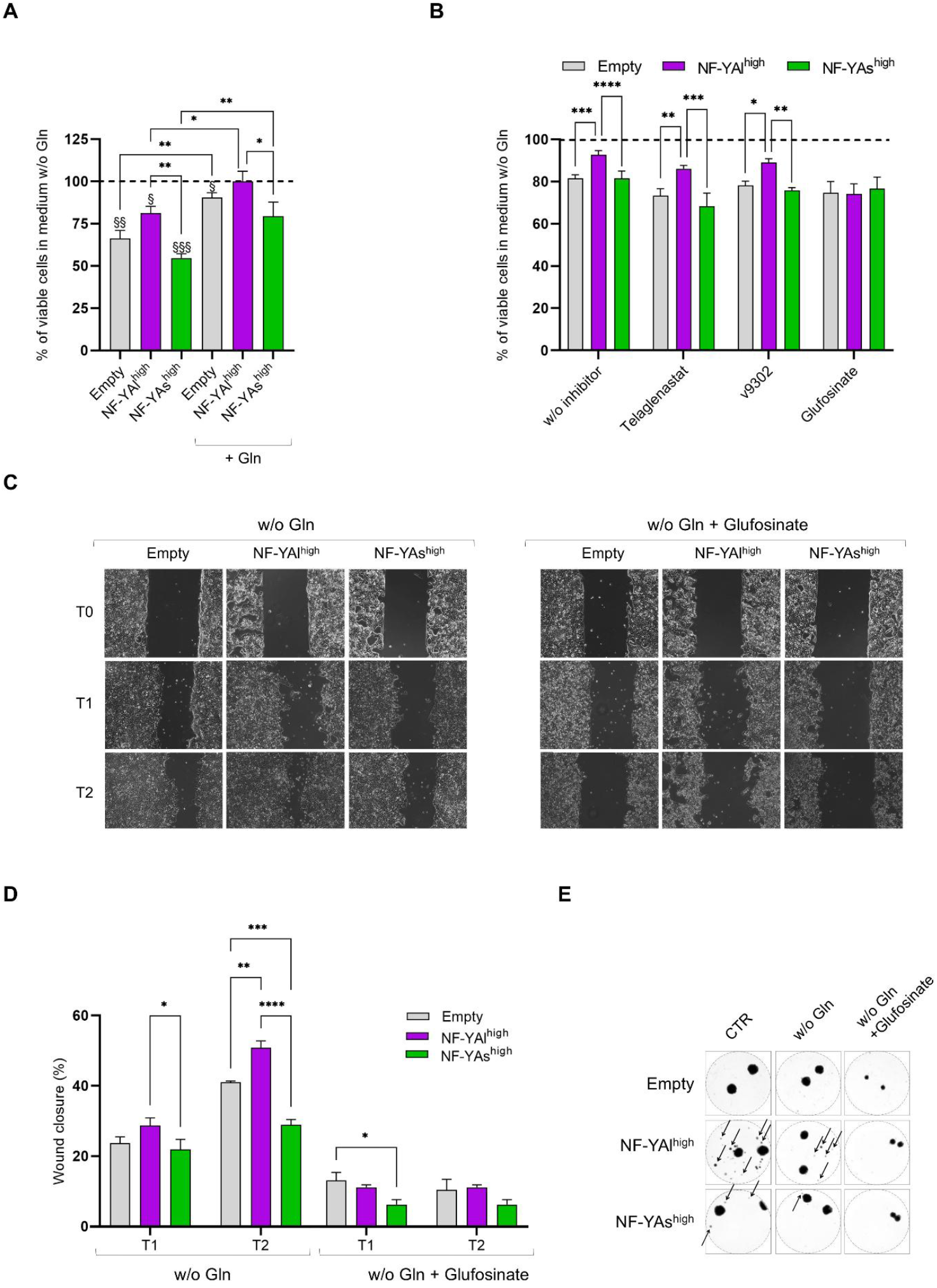
NF-YAl-driven glutamine synthesis supports proliferation and migration of CRC cells under glutamine-deprived conditions. **A)** Cell viability of Empty, NF-YAl^high^, and NF-YAs^high^ HCT116 cells analyzed by PrestoBlue cell viability assay after 72h in Gln-deprived medium or in Gln-deprived medium + 2 mM Glutamine. Data were normalized to cell growth in complete CTR medium, arbitrarily set at 100% (dotted line), and represent mean±SEM (one-sample t-test for comparison with control medium (§); one-way ANOVA with Fisher’s LSD test for comparison between samples (∗); n=4). **B)** Cell viability of Empty, NF-YAl^high^, and NF-YAs^high^ HCT116 cells analyzed by PrestoBlue cell viability assay after 72h in Gln-deprived medium either w/o or with inhibitors (Glufosinate, V-9302 or Telaglenastat). Data were normalized to cell growth in complete CTR medium, arbitrarily set at 100% (dotted line), and presented as mean±SEM (two-way ANOVA with Fisher’s LSD test for comparison between samples; 3≤n≤6). **C)** Representative images of cell migration in wound-healing assay. Confluent cells were seeded into Ibidi wound-healing chambers and images were acquired at 0, 24h, and 48h after the culture chambers were removed. **D)** Quantification of migration in Ibidi wound-healing chambers is shown as wound closure (%) compared to the initial gap, arbitrarily set at 100%. Data represent mean ± SEM (two-way ANOVA with Fisher’s LSD test; n=3). **E)** Optical microscopy images representative of spheroid-based metastatic potential assay. Two spheroids generated from Empty, NF-YAl^high^, and NF-YAs^high^ HCT116 were transferred into tissue-culture 48-well plates and then grown in complete IMDM medium (CTR) or medium without Gln ± 10 μg/ml Glufosinate. Images were taken after 10 days to determine whether the cells were able to move from spheroids and establish colonies. Arrows indicate representative colonies.

To better understand the mechanism underlying resistance to Gln deprivation in NF-YAl^high^ cells, we inhibited key components of Gln metabolism under Gln-starvation conditions: Glutaminase by Telaglenastat, the transporter Slc1a5 by V-9302, and GS by Glufosinate. Comparison across the three isogenic cell lines showed that only GS inhibition abolished the previously observed differences in HCT116 and SW480 cells (Fig. 6B, Suppl. Fig. 3D). These results suggest that the increased resistance, and potentially aggressiveness, of CRC cells with high NF-YAl levels is driven by enhanced ability to synthesize Gln, rather than uptake or catabolism.

To test this hypothesis, we quantified intracellular Gln and found that it is lower in NF-YAl^high^ and in NF-YAs^high^ cells compared to Empty cells, although the difference was not statistically significant (Suppl. Fig. 4A). Measurements of extracellular Gln revealed higher Gln in the media of both NF-YAl^high^ and NF-YAs^high^ cells, suggesting that reduced uptake or increased export could contribute to the observed Gln differences (Suppl. Fig. 4A). Under Gln-deprivation, NF-YAl^high^ cells maintained higher extracellular Gln compared to NF-YAs^high^ or Empty cells, indicating active endogenous Gln synthesis and export (Suppl. Fig. 4B), consistent with the observed changes in *Glul* gene expression. Together, these data support a model in which NF-YAl promotes metabolic adaptation through enhanced glutamine biosynthesis, conferring resistance to nutrient stress.

We recently demonstrated that increased cell migration represents one of the main traits of NF-YAl^high^ cells aggressiveness [17]. Therefore, we determined whether targeting Gln metabolism can affect the migratory phenotype. Increased migration ability was observed in NF-YAl^high^ cells compared to both control and NF-YAs^high^ cells also in Gln-depleted medium (Fig. 4C), while the GS inhibitor strikingly arrested cell migration in all cell types, aggressive NF-YAl^high^ cells included (Fig. 4C, D).

The impact of NF-YAl levels on metastatic potential was further evaluated using a tumor spheroid-based migration assay, which recapitulates tumor cell dissemination from a solid tumor. Multicellular spheroids of HCT116 cells were formed using the hanging drop method and then transferred to culture plates to assess their ability to disseminate and form visible colonies (Fig. 4E), which reflect both migration and clonogenic growth efficiency of tumor cells. Although reduced in Gln-depleted conditions, NF-YAl^high^ cells generated more secondary colonies than control or NF-YAs^high^ cells. This effect was mitigated by treatment with the GS inhibitor Glufosinate, suggesting that NF-YAl^high^-driven metastatic potential depends on intracellular Gln synthesis.

### 2.5 Glutamine metabolic rewiring by NF-YAl enhances CRC stress resistance and *in vivo*

**metastasis**

When cancer cells detach from the primary tumor and enter the bloodstream as circulating tumor cells (CTCs), they are exposed to hemodynamic shear stress, which induces oxidative stress and apoptotic cell death. Concurrently, CTCs undergo profound metabolic rewiring to cope with the changes in nutrient availability and redox balance. Only those CTCs capable of withstanding these mechanical and metabolic challenges acquire enhanced metastatic potential [27]. Gln metabolism supports CRC cell survival in circulation by sustaining glutathione (GSH) levels, mitigating oxidative damage, and increasing their chances of colonizing distant sites. Consistently, NF-YAl^high^ cells, which display enhanced metastatic abilities as previously demonstrated [17], exhibit significantly higher GSH levels compared to empty and NF-YAs^high^ cells, further supporting a functional link between NF-YAl high expression, redox homeostasis, and metastatic fitness (Fig. 5A).

Gln levels in serum are associated with poorer overall survival and progression-free survival in CRC patients [32]. Therefore, we investigated whether high NF-YAl expression provides protection against oxidative stress under low-Gln conditions. NF-YAl^high^ cells exhibit lower superoxide levels under Gln deprivation and even upon H_2_O_2_ exposure, consistent with a Gln-driven redox program that may underpin their increased metastatic potential (Fig. 5B).

**Figure 5.**
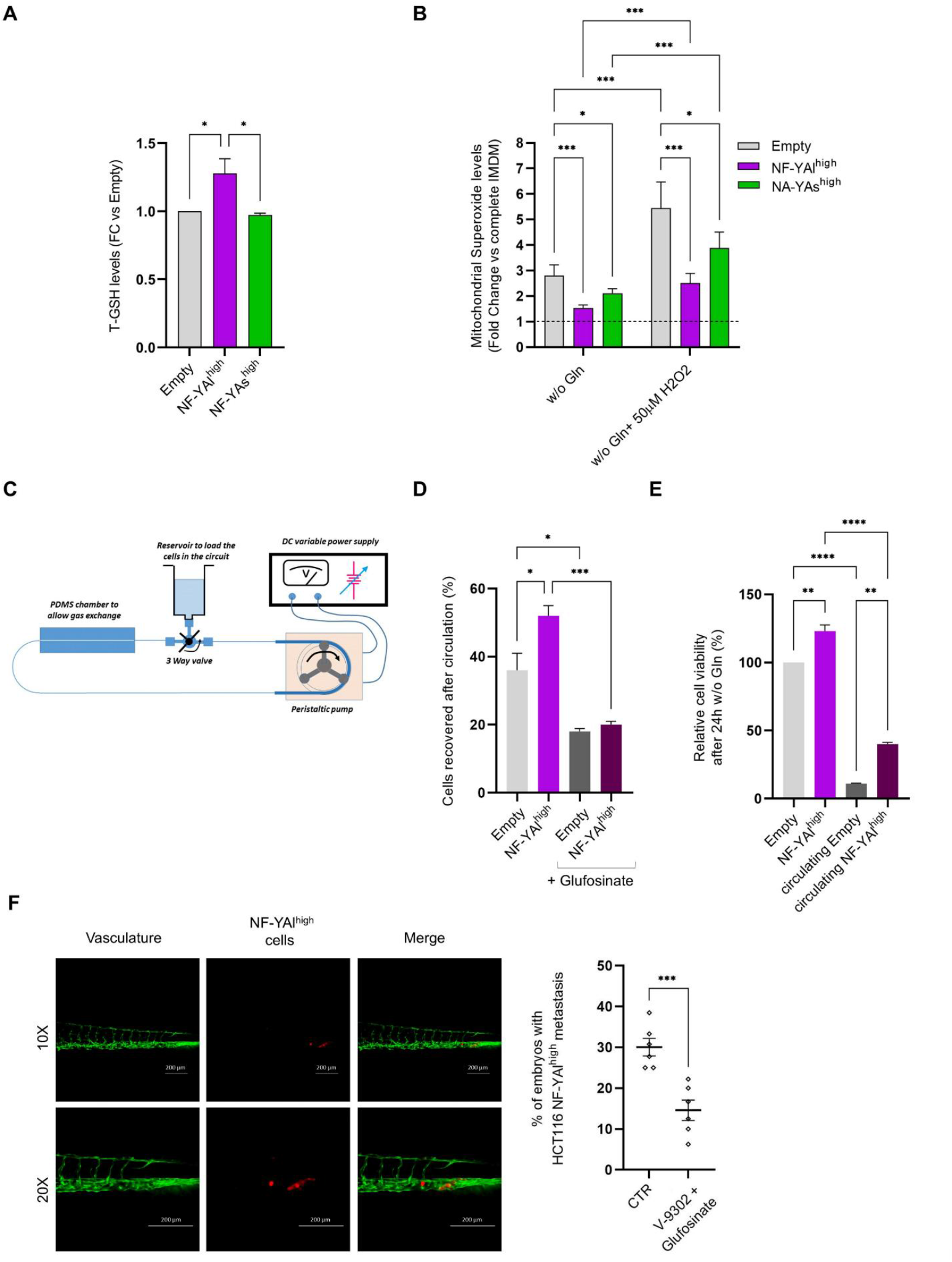
NF-YAl-dependent glutamine metabolic rewiring confers resistance to mitochondrial and shear stress under glutamine-depleted conditions and promotes CRC metastatization *in vivo*. **A)** Total GSH levels measured in NF-YAl^high^ and NF-YAs^high^ cells compared with Empty cells. Data are reported as fold change relative to Empty cells, arbitrarily set at 1. Data represent mean ± SEM (one-way ANOVA with Fisher’s LSD test; n=3). **B)** Mitochondrial superoxide production quantified with the MitoSox red dye in Empty, NF-YAl^high^, and NF-YAs^high^ HCT116. Cells were incubated for 72h in complete IMDM medium or in medium without Gln, and with or without the addition of 50µM H2O2. Data were normalized to cell growth in complete medium, arbitrarily set at 100% (dotted line), and represented as mean±SEM (two-way ANOVA with Fisher’s LSD test; n=4). **C)** Representative illustration of the microfluidic system developed for the analysis of cell survival under shear stress conditions, to mimic cell dissemination through body vessels. **D)** HCT116 Empty and NF-YAl^high^ cells were counted after 1h of circulation in Gln-free IMDM, with or without the addition of Glufosinate. Data are presented as the number of recovered cells (%) compared to the initial pre-circulation cell count, arbitrarily set at 100%. Data represent mean ± SEM (one-way ANOVA with Fisher’s LSD test; n=3). **E)** Equal amounts (500µl) of initial pre-circulation HCT116 Empty and NF-YAl^high^ cell suspension and of cells after 1h of circulation were plated in Gln-free IMDM in standard tissue culture plates. After 24h, cell number was determined. Data are presented as the number of cells (%) compared to HCT116 Empty cells, arbitrarily set at 100%. Data represent mean ± SEM (one-way ANOVA with Fisher’s LSD test; n=2). **F)** Left panel: representative immunofluorescence images of cell dissemination of NF-YAl^high^ HCT116 cells (red) 24h after injection in Tg(fli1: EGFP) zebrafish embryos (green). Scale bar: 200 μm. Right panel: Quantification of the number of embryos showing metastatic dissemination of NF-YAl^high^ cells in the presence of the metabolic inhibitors V-9302 and Glufosinate, compared with untreated controls. For treatment condition, before injection cells were pre-treated for 2h in Gln-free medium with 10 μg/ml Glufosinate and 1µM V-9302. Data represent mean ± SEM (t-test, n=6).

We next asked whether the metabolic adaptations of NF-YAl^high^ cells enhance resistance to circulatory stress, mimicking hemodynamic shear in a flow circuit model (Fig. 5C). NF-YAl^high^ cells showed higher viability after 1 hour of circulation in Gln-free medium, a survival advantage dependent on GS activity, as Glufosinate reduced recovered cells (Fig. 5D). This advantage persisted, with more NF-YAl^high^ cells capable of proliferating in Gln-free medium 24 hours post-circulation compared with controls (Fig. 5E).

Finally, we analyzed *in vivo* migration, building on our previous observation that NF-YAl^high^ cells exhibit increased metastatic potential in zebrafish xenografts [17]. To investigate the role of Gln metabolism *via* GS in the survival and dissemination of circulating CRCs, we pretreated NF-YAl^high^ cells to inhibit Gln uptake (V-9302) and synthesis (Glufosinate) before injection into the perivitelline space of zebrafish embryos. Treatment significantly reduced the embryos with metastatic dissemination, indicating that Gln metabolic rewiring is essential for NF-YAl^high^-driven metastasis *in vivo* (Fig. 5F).

### 2.6 NF-YAl–driven glutamine metabolism shapes the tumor microenvironment and modulates immune crosstalk in CRC patients

Analysis of CRC tumors from TCGA dataset revealed a weak positive correlation between NF-YAl and *Glul* expression in the overall cohort (R = 0.268), with a stronger relationship in CMS4 tumors (R = 0.396), where NF-YAl is highly expressed and linked to aggressive tumor characteristics [17] (Suppl. Fig. 4C). Kaplan-Meier analysis showed that only combined high NF-YAl and high *Glul* expression predicts poorer survival (Fig. 6A), suggesting that NF-YAl is required to support *Glul* adverse prognostic impact and may drive metabolic rewiring in aggressive CMS4 CRC.

**Figure 6.**
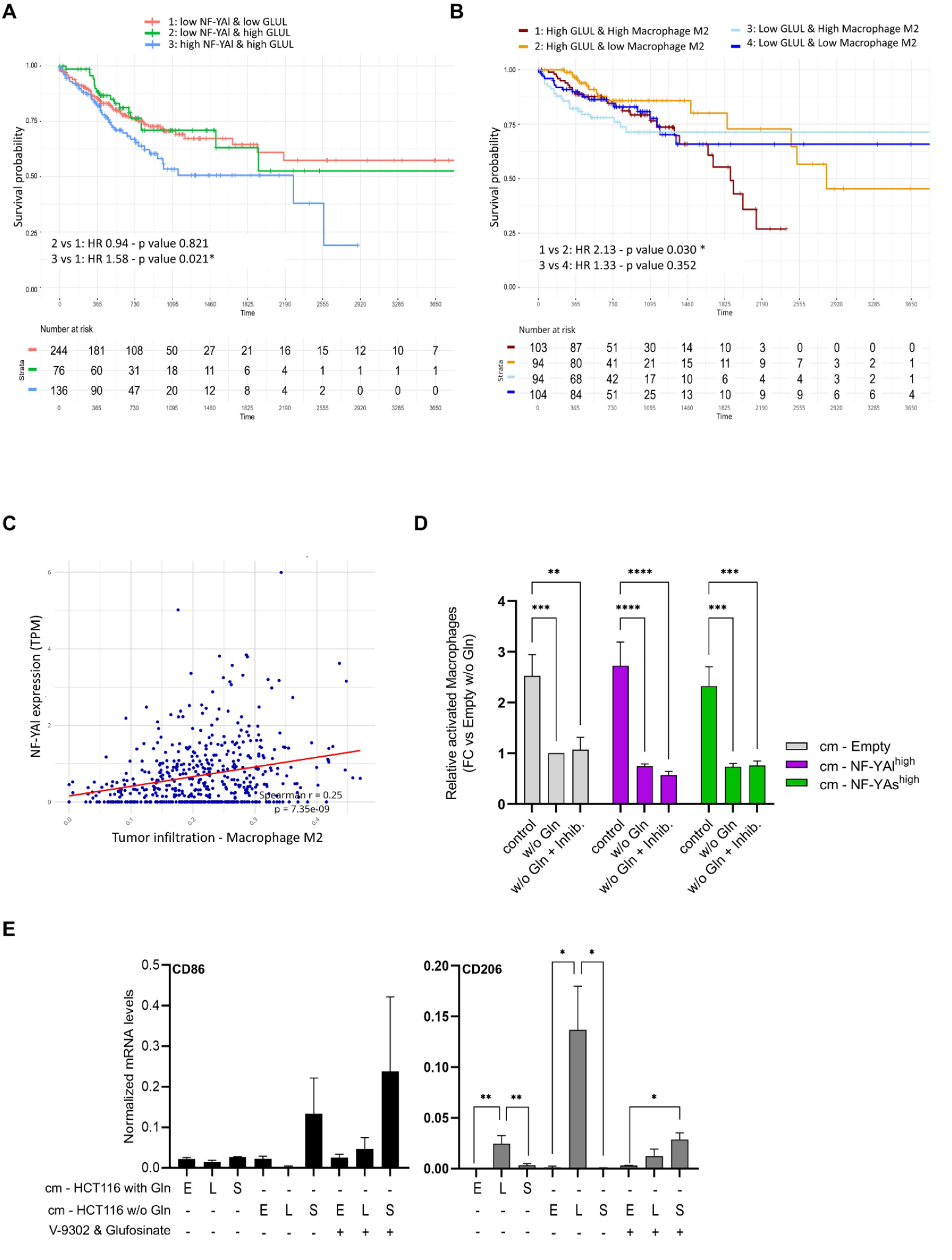
Glutamine metabolism in NF-YAl^high^ cancer cells of CRC patients shapes the tumor microenvironment and predicts survival probability. **A)** Kaplan–Meier analysis of survival probability measured in TCGA-COAD patients stratified according to high and low expression of NF-YAl together with high or low levels of *Glul*. Hazard ratio analyses (HR) are reported. **B)** Kaplan–Meier analysis of survival probability of TCGA-COAD patients stratified according to high and low expression of NF-YAl together with high or low levels of M2 macrophages infiltration. Hazard ratio analyses (HR) are reported. **C)** Correlation analysis between tumor-infiltration of M2 macrophages and the tumour mRNA levels of NF-YAl in TCGA-COAD patients. P-value and Spearman correlation coefficient are indicated. **D)** Human THP-1 monocytes were differentiated into macrophages by incubation in 50% (v/v) conditioned-medium (cm) of stable HCT116 cells grown in complete IMDM (control) or Gln-deprived IMDM. Glufosinate and V-9302 were added in the differentiation medium of macrophages when indicated. Differentiated macrophages were quantified by PrestoBlue assay after 48h. Data were normalized to THP-1 activation by cm of HCT116 Empty cells grown without Gln, arbitrarily set at 1. Data represent mean±SEM (two-way ANOVA with Fisher’s LSD test; n=4). **E)** THP-1 monocytes were induced to differentiate as described above, before RT-qPCR analysis. Transcript levels of CD68 (M1 marker) and CD206 (M2 marker) were analysed to verify their polarization. Results are reported as normalized mRNA levels and Rps20 was used as reference gene. Data represent mean±SEM (one-way ANOVA with Fisher’s LSD test; n=3).

CMS4 tumors are enriched in pro-tumorigenic M2 Tumor Associated Macrophages (TAM). High Gln consumption and glutaminolysis in TAMs are associated with M2 polarization, suggesting a link between NF-YAl-dependent metabolism of CRCs and TAM-mediated tumor support [28,29]. Based on these observations, we analyzed the TCGA-COAD dataset using the CIBERSORT deconvolution algorithm [30]. Tumors with high *Glul* expression and abundant M2 macrophage infiltration showed poorer overall survival (Hazard Ratio 2.13) compared to those with high *Glul* but low M2 infiltration (Fig. 6B), suggesting that *Glul* expression plays a critical role in shaping the TME. We therefore examined the relationship between NF-YAl levels and TAMs, finding a significant positive correlation between NF-YAl levels, but not NF-YAs, and M2 macrophages infiltration (Fig. 6C, Suppl. Fig. 4D).

To assess Gln-based metabolic symbiosis between NF-YAl^high^ cells and macrophages, THP-1 monocytes were exposed to conditioned medium (CM) from HCT116 cells cultured with or without Gln [31]. Gln depletion reduced monocyte-to-macrophage differentiation by ~2.5-fold across CM from Empty, NF-YAl^high^ and NF-YAs^high^ cells, indicating reliance on exogenous Gln (Fig. 6D) [32]. Inhibiting endogenous Gln synthesis and uptake did not further affect their activation. Despite this, RT-qPCR of macrophage markers revealed distinct polarization patterns: CM from NF-YAs^high^ increased CD86 (M1), while CM from NF-YAl^high^ strongly induced CD206 (M2), indicating pro-tumoral TAM activation. Notably, blocking Gln uptake in THP-1 cells reversed M2 polarization, highlighting the pivotal role of NF-YAl-driven Gln production in modulating the TME.

## 3. Discussion

Recent transcriptomic data from primary and metastatic CRCs highlighted an enrichment of the GO term “Gln metabolism” in both paired and non-paired metastatic lesions *versus* primary tissues [33]. Despite this, low levels of Gln in the serum of CRC patients correlate with high systemic inflammation and poor prognosis [34]. Expression of the glutamate ammonia ligase gene (*Glul*) is significantly higher in metastatic tumors than in primary lesions [33], underscoring the capacity of metastatic cells to adapt to the hostile TME by increasing endogenous Gln production, which enhances survival and resistance to oxidative stress and nutrient scarcity [27]. Based on the findings that GLS1^KD^ promotes apoptosis and reverses oxaliplatin resistance of CRC cells [35,36], pharmacological strategies aimed at inhibiting GLS have been explored [37]. However, the capacity of tumor cells to produce Gln endogenously may limit the efficacy of such interventions, emphasizing the need for a deeper investigation into both catabolic and anabolic pathways involved in Gln metabolism.

Building on the demonstrated transcriptional activity of NF-Y in metabolic pathways altered in CRC cells [11], we investigated the metabolic profile of CRC cells depleted of the NF-YA regulatory subunit and found halved levels of Glu in NF-YA^KD^ cells. On the other hand, NF-YAl^high^ cells, characterized by mesenchymal phenotype and enhanced metastatic potential *in vitro* and *in vivo* [17], also display distinct Gln metabolic features. Specifically, NF-YAl^high^ cells are more resistant to Gln deprivation than control or NF-YAs^high^ cells, despite relying more heavily on Gln oxidation to sustain baseline mitochondrial respiration. This indicates that NF-YAl expression promotes metabolic adaptations that support cell survival and energy production under Gln-limiting conditions.

Multiple genes involved in Gln metabolism are altered by NF-YA modulation, with resistance to Gln deprivation being associated with the transcriptional up-regulation of the *Glul* gene in NF-YAl^high^ cells, in accordance with a previous study from Dolfini et al. in osteosarcoma cells [38]. Our deeper functional characterization and mechanistic analysis demonstrated that NF-Y binds to the *Glul* promoter, and increased NF-YAl recruitment impacts on chromatin accessibility, marked by higher H3K4 methylation. The question of whether NF-YAl has higher DNA affinity compared to NF-YAs, or if the exon 3-encoded peptide in the NF-YAl transactivation domain allows its interaction with different coactivators, including chromatin modifiers, remains to be explored.

NF-YAl^high^ cells preserve their increased invasive abilities even in the absence of Gln, a condition that mimics the reduced Gln levels of high-risk patients, as demonstrated by migration and colony assays and shear stress tests. Inhibition of Gln synthesis through Glufosinate administration significantly reduces the aggressiveness of NF-YAl^high^ cells, markedly impairing their migratory and metastatic potential both *in vitro* and *in vivo*, thereby revealing a critical metabolic vulnerability in CRC tumors with high NF-YA expression (CMS4 subtype) [17].

CMS4 tumors undergo extensive TME remodeling that enforces EMT programs and drives poor clinical outcomes. A central component of this remodeling is metabolic rewiring: CRC cells shift glucose, glutamine/tryptophan, and fatty acid metabolism in ways that suppress anti-tumor immunity and accelerate tumor progression [39]. Within the TME, Gln availability directs macrophage polarization toward an M2/TAM phenotype, thereby supporting tumor growth, angiogenesis, dissemination, and therapeutic resistance [40]. Our findings identify NF-YAl, but not NF-YAs, as a key regulator of the metabolic crosstalk between CRC cells and the TME, specifically influencing TAM plasticity through Gln metabolism. Within the CMS4 subtype, NF-YAl shows a strong correlation with Glul expression, supporting a model in which NF-YAl enhances tumor Gln synthesis and release. This metabolic rewiring is associated with increased infiltration of M2-like macrophages in NF-YAl^high^ tumors. Functional experiments using conditioned media of CRC cells further support this model, demonstrating that macrophage activation and maintenance of an M2-like phenotype are highly dependent on Gln release *via* a NF-YAl/*Glul* axis. Inhibition of Gln utilization in macrophages reverses the M2 program induced by NF-YAl^high^ conditioned media.

These findings reveal a direct link between NF-YAl-driven Gln metabolism and TAM reprogramming, sustaining pro-tumorigenic states in CMS4 CRC. CMS4 tumors, with high NF-YAl expression and abundant M2-like TAMs, may be particularly vulnerable to therapies targeting Gln production from cancer cells, simultaneously limiting tumor growth, metastasis, and TAM-mediated support (graphical abstract).

**Graphical Abstract**

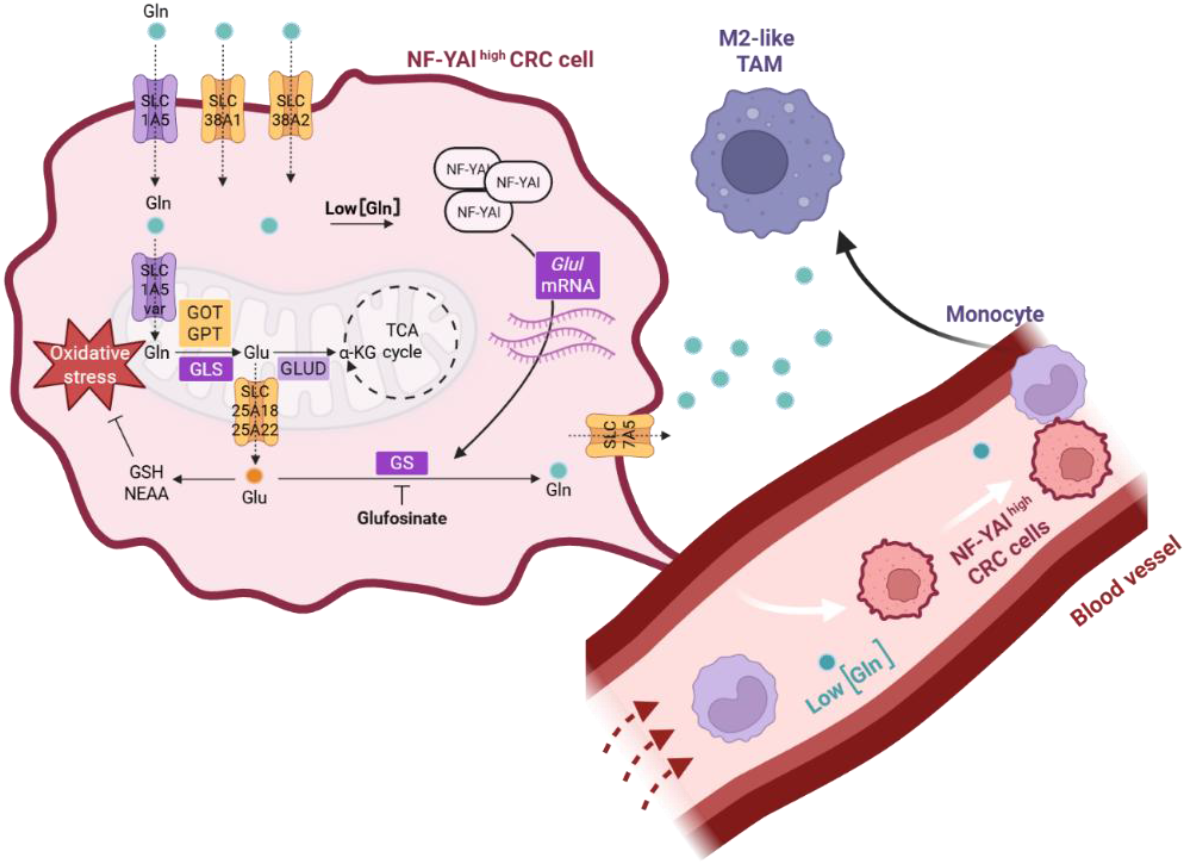

## 4. Material and Methods

Detailed materials and methods are supplied as Supplementary Methods.

### TCGA data analysis

TCGA data analysis exploited data from the Colorectal Adenocarcinoma dataset, integrating Illumina Hiseq and GA data from FireBrowse (http://firebrowse.org/) with clinical and survival data from UCSC Xena Browser [41]. CMS-based classification was retrieved from published data [17]. Estimated tumor infiltration data were retrieved from TIMER 2.0 [42] selecting CIBERSORT for downstream analyses.

Statistical analyses and visualizations were performed in R environment v4.4.3, using the *tidyverse* suite [43], *finalfit* and *ComplexHeatmap* [44]. NF-YAl and GLUL expression clusters were identified by k-means clustering (k=4, seed=3000) applied to the expression matrix in log2(TPM+1) after z-score normalization. Correlations were computed using cor.test function with spearman method.

### Cell lines and lentiviral transduction

Human colon cancer HCT116 and SW-480 cells, and monocytic THP-1 cells were grown in standard conditions. NF-YA^KD^ HCT116 [45] and stable NF-YA-overexpressing cells [17] were obtained by lentiviral infection of colon cancer cells as previously reported.

### Glutamine starvation and cell treatments

Gln depletion experiments were performed in Gln-free medium, supplemented with 10% Dialyzed FBS (HyClone™ dialyzed Fetal Bovine Serum, #SH30079). When indicated, we concomitantly added Gln pathways inhibitors: 10 μg/ml Glufosinate-ammonium (Sigma-Aldrich, #45520), 1µM V-9302 (MedChemExpress, #HY-112683A) or 25nM Telaglenastat -CB-839-(MedChemExpress, #HY-12248).

### Cell proliferation assay

Cell viability was evaluated by PrestoBlue reagent (#A13261, Thermo Fisher Scientific, MA), according to the manufacturer’s protocol.

### 2D and 3D cell migration assays

Wound healing assays were performed with *Ibidi* culture-inserts (#80209, Ibidi GmbH, Germany) as previously described [17].

For micrometastasis assay, two Multicellular Tumor Spheroids (MTSs) generated from HCT116 cells were transferred into 48-well plates and grown in complete IMDM medium or medium w/o Gln ± 10 μg/ml Glufosinate for 10 days. MTSs and MTS-derived colonies were fixed and stained with 0.1% crystal violet solution.

Plates were imaged by EVOS M5000 imaging system (Thermo Fisher Scientific, MA).

### Protein extraction and Immunoblotting

Cells were lysed into 1X SDS sample buffer [17] and western blots were performed with nitrocellulose membrane in a Trans-Blot Turbo Transfer System (Bio-Rad, USA). Primary and secondary antibodies are indicated in Supplementary Methods.

### RT-qPCR

RealTime PCRs were performed with SsoAdvanced Universal SYBR Green Supermix (#1725274, Bio-Rad, USA) using Biorad CFX connect Real-Time PCR Detection System. Oligonucleotides sequences are listed in Suppl. Table 3. Control cell line or control growth medium was used as reference sample, as indicated in figure legends.

### Promoter analysis

In silico analysis of NF–Y motifs in promoters (−950 to +50 relative to TSS) was performed with the computational algorithm Lasagna-Search 2.0 [24]. The presence of the CCAAT/ATTGG sequence and of NF–Y binding from the ENCODE Project was manually assessed in the UCSC Genome Browser (human GRCh38/hg38).

### Chromatin immunoprecipitation (ChIP)

ChIP was performed as previously described [46] with 1 µl of anti-NF-YA (#C15310261, Diagenode), 2 µg of anti-H3K4me3 (#C15410003, Diagenode) or control IgG (#30000-0-AP, Proteintech). Results are presented as % of INPUT DNA. Oligonucleotides sequences are listed in Suppl. Table 3.

### Zebrafish Xenograft Injection of cancer cells

Adult zebrafish and Tg(fli1a:EGFP) embryos were maintained per the European Directive 2010/63/EU and Italian law (D.Lgs. 26/2014). NF-YAl^high^ HCT116 cells were pre-treated for 2h in Gln-free medium ± 10 μg/ml Glufosinate-ammonium and 1µM V-9302. Cells were labelled with CellTracker Orange CMRA viable dye (Invitrogen, Carlsbad, CA, USA), resuspended in PBS (CTR) or treatment solution at 2.5 × 10^5^ cells/µL, and grafted into the perivitelline space of 48 hours post-fertilization Tg(fli1a:EGFP) embryos. Circulating cells were imaged in vivo after 24h using a fluorescence stereomicroscope (Nikon SMZ25) and analyzed with NIS-Elements software (Nikon Corporation, Tokyo, Japan).

### *ex vivo* High Resolution Magic Angle Spinning (HR-MAS) NMR

HCT116 were harvested 48h-post infection with scramble (CTR) or NF-YA shRNA (NF-YA^KD^) lentiviral vectors. *Ex vivo* NMR analysis was performed using 4-6 x 10^6^ cells, acquiring one- and two-dimensional spectra [47]. Metabolite areas were estimated in 1D 1H CPMG spectra using Mnova software (MestReNova, ver. 8.1.0) [48], with relative concentration defined as the ratio of metabolite resonance area to cell numbers (arbitrary units). Figure S1 reports a representative spectrum. Metabolite set enrichment and integrated joint pathway analyses were performed using MetaboAnalyst 6.0 [49], analyzing differential metabolites with our matched transcriptomic dataset from NF-YA^KD^ HCT116 cells (GSE70543) or an independent siRNA-NF-YA dataset (GSE56788).

### Metabolic analysis using seahorse technology

Metabolic characterization of HCT116 cells was performed using a Seahorse XFe24 Extracellular Flux Analyzer (Seahorse Bioscience). Data were normalized to cell numbers. The Mito Fuel Flex Test measured the OCR and the dependency, capacity and flexibility of cells to oxidize glucose, glutamine or long-chain fatty acids.

### Detection of mitochondrial superoxide (MitoSOX)

5×10^3^ HCT116 cells were seeded in 96-well plate for 24h, then incubated in Gln-deprived medium ± 50µM H2O2 for 72h, and stained with the MitoSox red dye (Thermo Fisher Scientific, Inc.) according to the manufacturer’s procedures. Fluorescence was measured after 30 min using a GloMax Discover microplate reader (Promega).

### Measurement of glutamine and GSH

HCT116 cells were seeded at 1 x 10^4^/well in a 96-well plate for 24h, then treated with complete or Gln-depleted medium for another 24h. Gln was measured in cell media using the Glutamine/Glutamate-Glo Assay kit (Promega, Cat#: J8021), while total Glutathione was quantified into cells with a colorimetric kit (elabscience, Cat#: E-BC-K097-M), according to the manufacturer’s instructions. Measurements were taken using a GloMax Discover microplate reader (Promega).

### Microfluidic circulatory system fabrication and circulation of HCT116 colon cancer cells

We developed a circulating system exploiting a peristaltic pump and microfluidic devices. HCT116 cells at ~75% confluence were pre-treated for 1h in Gln-free IMDM, collected and suspended in Gln-free IMDM ± 10 µg/ml Glufosinate. 1.5 mL of HCT116 cell suspension (5 × 10^5^ cells/ml) was injected into the system and recovered after 1h. Cell number was determined with a NucleoCounter NC-100 (ChemoMetec). Post-circulation, 500µl cell suspension was plated in Gln-free IMDM for 24h before determining cell numbers. The initial pre-circulation suspension was used as control in recovery experiments.

### THP-1 monocytes differentiation into macrophages

THP-1 were incubated for 48h in a 50% mixture of filtered conditioned medium (CM) from HCT116 cells and RPMI with either complete or Gln-deprived supplements. Where indicated, 10 μg/ml Glufosinate and 1µM V-9302 were added in the Gln-depleted medium. CM was obtained by seeding 1.5 x 10^5^ HCT116 in 6-wells plates, replacing the medium after 18h, and collecting the CM after 24h. Activated monocytes became adherent and we assessed cell viability and macrophage markers by RT-qPCR.

### Statistical analysis

All graphs represent the mean of at least three independent experiments ± SEM. Statistical analyses were performed using GraphPad PRISM software using ANOVA or Student’s t-test, as appropriate. p values of p < 0.05 were considered statistically significant: p<0.05(*), p<0.01(**), p<0.001(***), and p<0.0001(****).

## Abbreviations

Colorectal cancer (CRC), Tumor Microenvironment (TME), Epitelial-to-Mesenchymal Transition (EMT), Conditioned Medium (CM), Extracellular Matrix (ECM), Tumor Microenvironment (TME), Tumor Associated Macrophage (TAM), Alanine (Ala), Acetate (Ac),N-Acetyl-Aspartate (NAA), Branched-Chain Aminoacid (BCAA), Creatine (Cr), Glutamine (Gln), Glutamate (Glu), Glutathione (GSH), Glycine (Gly), Histidine (Hys), Isoleucine (Ile), Lactate (Lac), Leucine (Leu), Phenylalanine (Phe), Phosphocholine (PC), Piruvate (Pir), Taurine (Tau),Threonine (Thr), Valine (Val)

## Acknowledgements

We thank Dr. Cristina Ruberti at the University of Milan for the valuable technical support in Seahorse cell metabolism experiments.

## Author contributions

**C.I**.: Conceptualization, Writing – original draft, Supervision, Project administration, Funding acquisition. **S.B**.: Conceptualization, Investigation, Methodology, Formal analysis, Visualization, Writing – original draft. **V.M**.: Investigation, Methodology, Formal analysis. **V.R**., **M.R**., **G.M**., **O.C**.: Investigation, Methodology. **N.I**., **V.C**., **L.C**., **A.R, T.M**.: Investigation, Validation. **V.S**.: Investigation. **A.A**.: Methodology, Resources. **D.D**.: Formal analysis. **V.Z**.: Supervision, Funding acquisition. All authors read and approved the final manuscript.

## Funding

The research leading to these results has received funding from AIRC (Fondazione Italiana per la Ricerca sul Cancro) under IG 2018 – I.D. 21323 project – P.I. Carol Imbriano and from the Department of Life Sciences, University of Modena and Reggio Emilia, under FAR2023PD grant - P.I. Silvia Belluti and FAR2025PD grant - P.I. Carol Imbriano. The project is funded under the National Recovery and Resilience Plan (NRRP), Mission 4 Component 2 Investment 1.4 - MUR public notice n. 3138/2021 as modified by DD 3175/2021 funded by the European Union – NextGenerationEU, CN_3: National Center for Gene Therapy and Drugs based on RNA Technology Spoke 2 - Cancer”, project code CN 00000041 to C.I., and NRPP, Mission 4 Component 2 Investment 1.3 - MUR public notice n. 341 of 03.15.2022 funded by the European Union – NextGenerationEU, Creation of “Partnerships extended to universities, research centers, companies for the financing of basic research projects”, HEAL ITALIA Spoke 3, project code PE00000019 to V.Z. The funders played no role in study design, data collection, analysis and interpretation of data, or the writing of this manuscript. Views and opinions expressed are however those of the authors only and do not necessarily reflect those of the European Community or European Union. Neither the European Union nor the European Community can be held responsible for them.

## Data availability

The data supporting the findings of this study are available within the article and its supplementary information files. Data of TCGA Colorectal Adenocarcinoma dataset are available from public repositories: RNA-seq data were retrieved from firebrowse.org merging Illumina Hiseq and GA data; Clinical and survival data were retrieved from UCSC Xena Browser (http://xena.ucsc.edu).

