## Supplementary information for "Metabolic reprogramming controlled by NF-YA alternative splicing creates therapeutic opportunities in colorectal cancer"

**SUPPLEMENTARY FILES**

a) **Supplementary FIGURES**
b) **Supplementary TABLES**
c) **Supplementary MATERIALS AND METHODS**

### a) Supplementary FIGURES

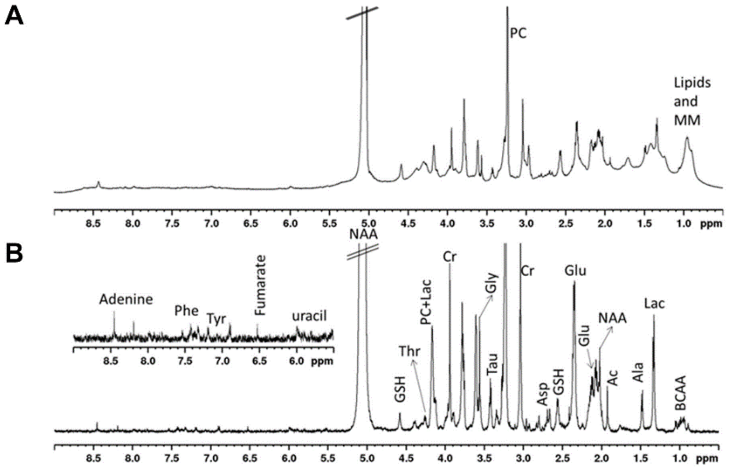


**Suppl. Figure 1. Representative spectra obtained by High Resolution Magic Angle Spinning (HR-MAS) NMR.** Representative A) $^{1}$H spectrum with water presaturation and B) cpmg spectrum. The visible and clear resonances are assigned. The major metabolites are labelled: creatine (Cr), lactate (Lac), taurine (Tau), macromolecules (MM), acetate (Ac), glutamate (Glu), glycine (Gly), (BCAA).

**A**

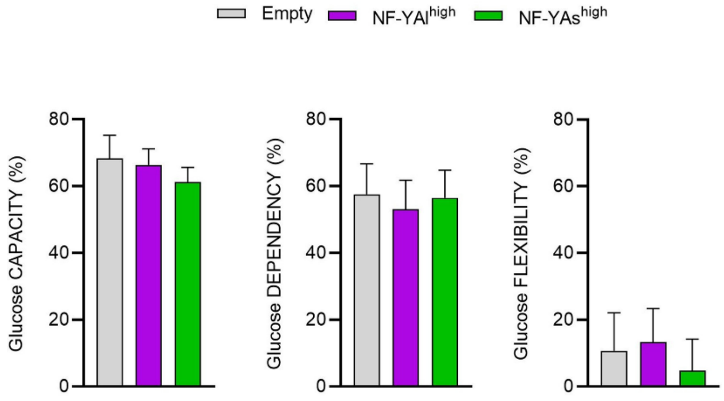


**B**

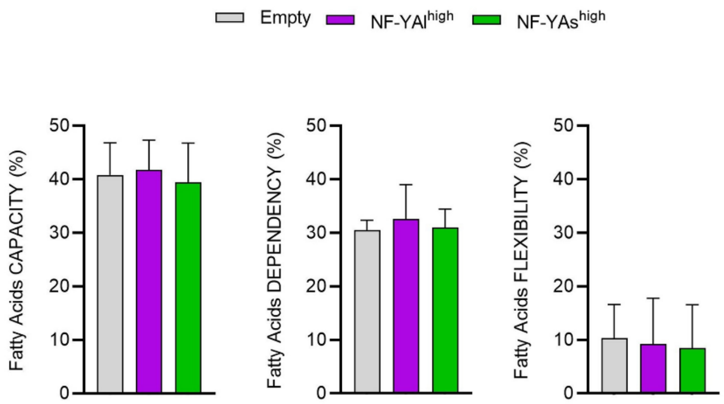


**Suppl. Figure 2. Identification of live metabolism in NF-YAl$^{high}$ and NF-YAs$^{high}$ cells using Seahorse analysis.** Effect of overexpression of NF-YAs and NF-YAl on A) glucose capacity, dependency and flexibility (%) and B) fatty acids capacity, dependency and flexibility (%). Data are expressed as mean±SEM (two-way ANOVA with Fisher's LSD test, n = 3).

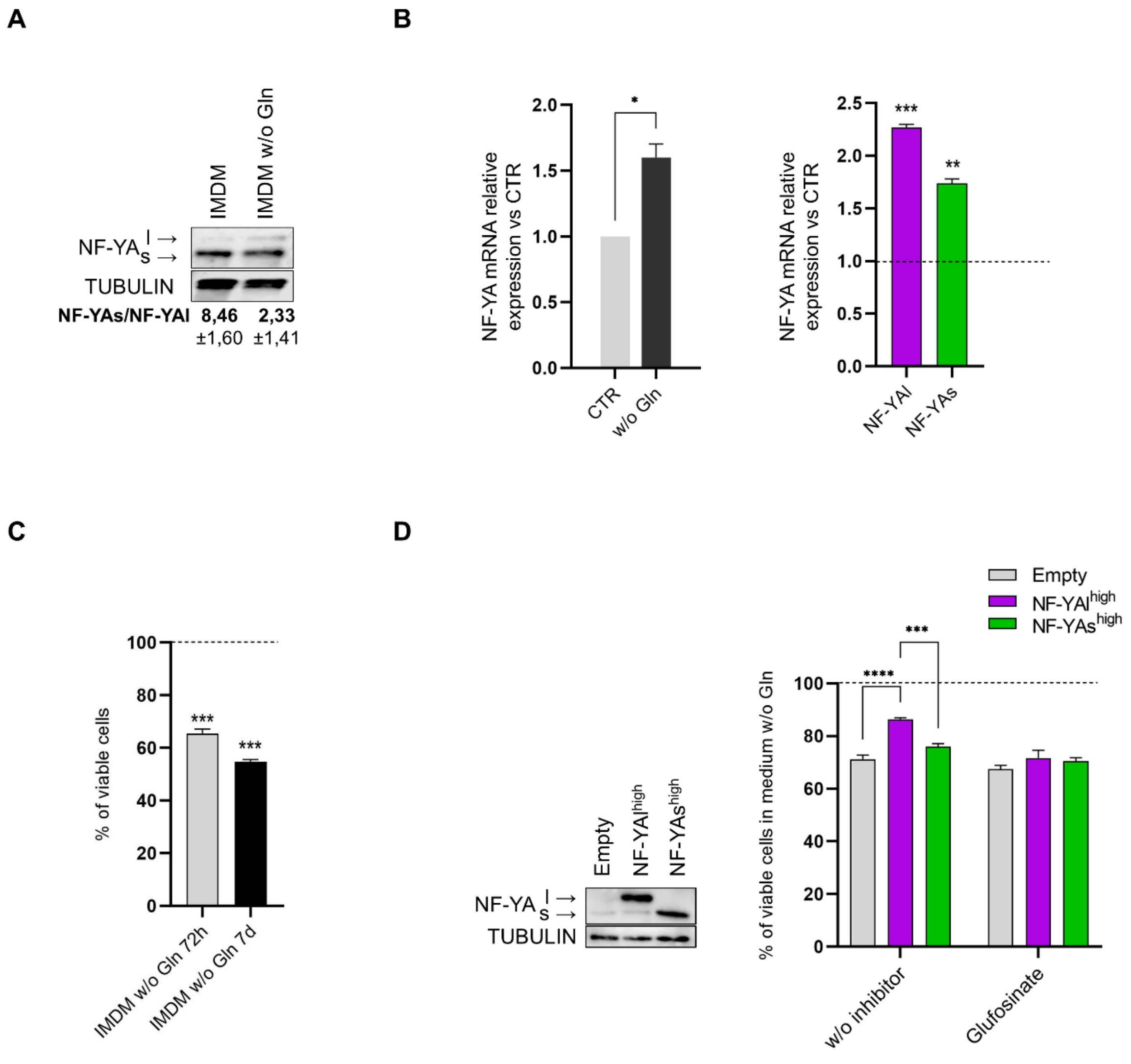


**Suppl. Figure 3. Effects of Glutamine deprivation and NF-YA isoforms expression in SW480 cells.** A) Protein levels of NF-YA measured by Western blot in SW480 cells in complete IMDM medium or IMDM without Gln. Tubulin was used as loading control. A representative quantification of NF-YAs and NF-YAl expression by ImageJ analysis and the NF–YAs/NF–YAl ratio is reported. B) Transcript levels of total NF-YA (left) and transcript levels of NF-YAl and NF-YAs (right) in complete medium (CTR) or w/o Gln, quantified by RT-qPCR. Results are reported as fold change vs CTR, arbitrarily set at 1 (dotted line in right panel). Rps20 was used as reference gene. Data represent mean±SEM (one- sample t-test; n=3). C) Cell viability of SW480 cells analyzed by PrestoBlue cell viability assay after 72h and 7 days in Gln-deprived medium. Data were normalized to cell growth in CTR medium, arbitrarily set at 100% (dotted line), and represent mean±SEM (one- sample t-test; n=3). D) Western blot analysis (left panel) of total extracts from SW480 cells stably infected with Empty, NF-YAl and NF-YAs lentiviral particles. Tubulin was used as loading control. Cell viability (right panel) of isogenic SW480 cells analysed by PrestoBlue cell viability assay after 72h in Gln-deprived medium either w/o inhibitors or in presence of Glufosinate. Data were normalized to cell growth in complete CTR medium, arbitrarily set at 100% (dotted line), and presented as mean±SEM (two-way ANOVA with Fisher’s LSD test; n=3).

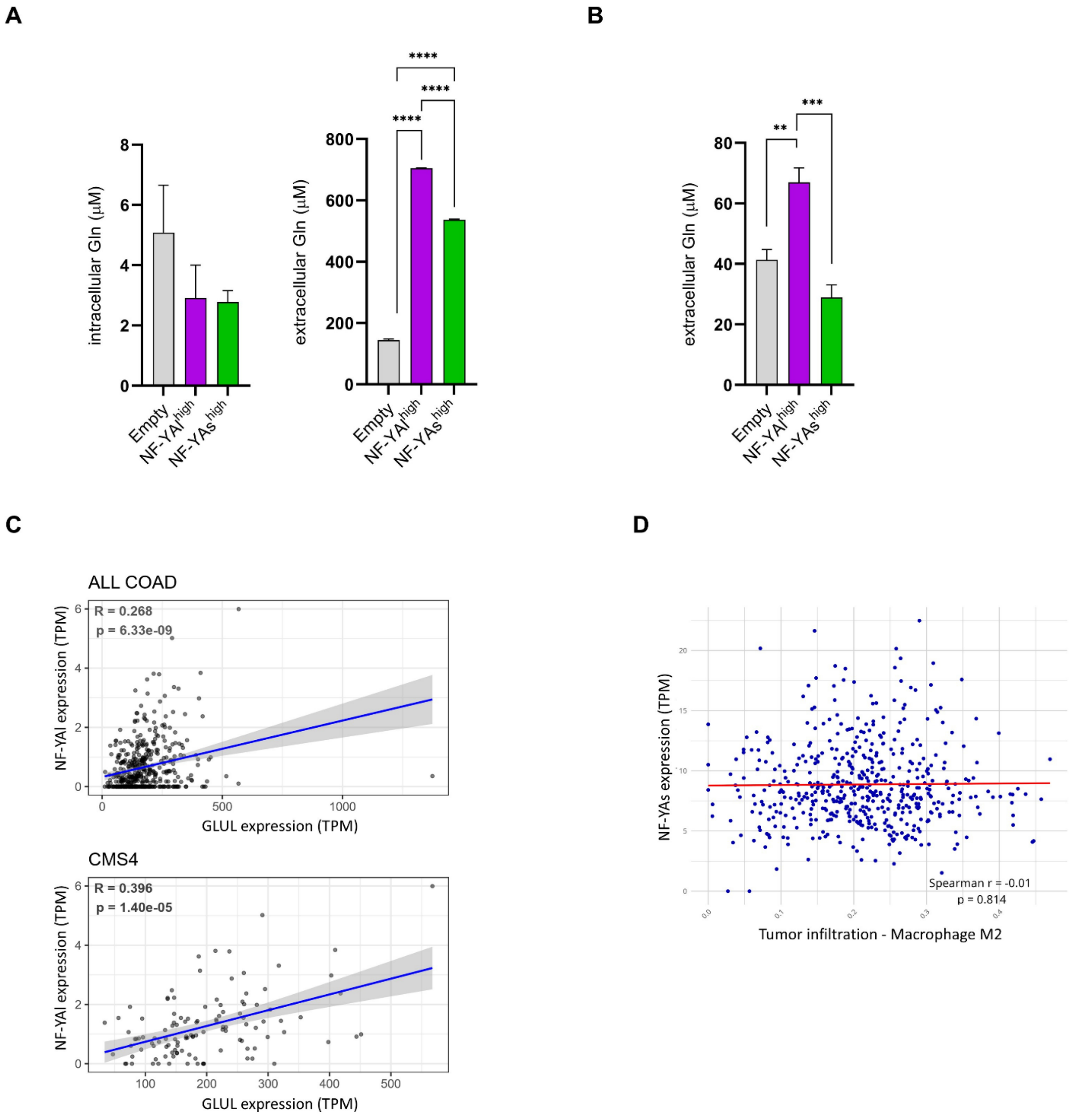


**Suppl. Figure 4. NF-YAl promotes enhanced glutamine biosynthesis and export, and it positively correlates with Glul levels in CMS4 tumors.** A) Left panel*:* Concentrations of intracellular glutamine measured in isogenic HCT116 cell lysates 24h after cells seeding in complete medium. Right panel: Concentrations of glutamine in growth media after 24h of cell growth in complete medium. Glutamine concentration was measured using the bioluminescent Glutamine/Glutamate-Glo Assay kit, according to manufacturer's instructions. Data represent mean±SEM (one-way ANOVA with Fisher's LSD test, n= 3 for media, n=4 for lysates). **B)** Extracellular concentrations of glutamine after 24h of cell growth of isogenic HCT116 cells in medium without glutamine. Data represent mean±SEM (one-way ANOVA with Fisher's LSD test, n= 3). **C)** Correlation analysis between NF-YAl and *Glul* transcript levels in all TCGA-COAD patients (upper panel) or specifically in CMS4 patients (lower panel). P-values and Spearman correlation coefficients are indicated. **D)** Correlation analysis between tumor-infiltration of M2 macrophages and the mRNA levels of NF-YAs in TCGA-COAD patients. P-value and Spearman correlation coefficient are indicated.

### b) Supplementary TABLES

**Suppl. Table 1.** List of $^1H$ and $^{13}C$ chemical Shift ($\delta$, ppm) of metabolites. [a],$^1H$ chemical shifts refer to Ala doublet at 1.48 ppm; [b].$^{13}C$ chemical shifts refer to Ala at 16.8 ppm; [c], contributes to the 3.77, 57.1 ppm cross-peak.

| | **Metabolites** | **$\delta^1H$ [a]** | **$\delta^{13}C$ [b]** | **group** |
|---|---|---|---|---|
| **1** | Fatty acids | 0.89 | 16.5 | $CH_3$ |
| | | 1.32-1.29 | 32.6-34.2 | $(CH_2)_n$ |
| | | 1.59 | 27.4 | $CH_2$-$CH_2$-C=O |
| | | 2.04 | 29.6 | $CH_2$CH=CH |
| | | 2.27 | 36.1 | $CH_2$-C=O |
| | | 2.78 | 27.9 | =CH-$CH_2$-CH= |
| | | 5.32 | 128-130 | -CH=CH- |
| **2** | Lactate | 1.33 (d) | 22.7 | $CH_3$ |
| | | 4.12 (q) | 71.3 | CH |
| **3** | Alanine | 1.48 (d) | 19.3 | $CH_3$ |
| | | 3.78 | c | $\alpha$-CH |
| **4** | Valine | 0.98 (d) | 21.7 | $\gamma$-$CH_3$ |
| | | 1.04 (d) | 20.7 | $\gamma$-$CH_3$ |
| | | 2.27 | 30.5 | $\beta$-CH |
| | | 3.61 | c | $\alpha$-CH |
| **5** | Leucine | 0.96 | 24.9 | $\delta$-$CH_3$ |
| | | 0.97 | 25.3 | $\delta$-$CH_3$ |
| | | 1.70 | 42.4 | $\beta$-$CH_2$ |
| | | 1.73 | 28.8 | $\gamma$-$CH_2$ |
| | | 3.72 | c | $\alpha$-CH |
| **6** | Lysine | 1.88 | 32.9 | $\beta$-$CH_2$ |
| | | 1.47 | 22.6 | $\gamma$-$CH_2$ |
| | | 1.73 | 27.3 | $\delta$-$CH_2$ |
| | | 3.03 | 39,7 | $\varepsilon$-$CH_2$ |
| | | 3.73 | c | $\alpha$-CH |
| **7** | Isoleucine | 1.97 (d) | 25.2 | $\gamma$-$CH_3$ |

| | | | | |
|---|---|---|---|---|
| | | 0.95-0.93 (t) | 21 | δ-$CH_3$ |
| **8** | Acetate | 1.92 (s) | 26.6 | $CH_3$ |
| **9** | Threonine | 1.32 | 21.8 | γ-$CH_3$ |
| | | 3.56 | 69 | β-CH |
| | | 4.25 | 66.9 | α-CH |
| **10** | Glutamate | 2.36 | 34.3 | γ-$CH_2$ |
| | | 2.07,2.13 | 25.2 | β-$CH_2$ |
| | | 3.73 | c | α-CH |
| **11** | Creatine | 3.06 (s) | 39.6 | $CH_3$ |
| | | 3.95 (s) | 56.4 | $CH_2$ |
| **12** | Glycine | 3.56 (s) | 44.3 | $CH_2$ |
| **13** | Taurine | 3.41 (t) | 38.06 | N-$CH_2$ |
| | | 3.26 (t) | 48.4 | S-$CH_2$ |
| **14** | Gluthatione | 2.55 | | 5 $CH_2$ |
| | | 2.95 | | 4 $CH_2$ |
| | | 4.57 | | 1 $CH_2$ |
| **15** | Histidine | 7.03 (s) | | 2 CH |
| | | 7.73 (s) | | 4 CH |
| **17** | Uridine | 5.89 5.93 | | CH |
| | | 7.91 | | CH |
| **18** | Ascorbate | 4.52 | | |
| **19** | Aspartate | 2.68-2.65 | | $CH_3$ |
| | | 3.90 | | $CH_2$ |
| **20** | Tyrosine | 6.88 | | |
| | | 7.18 | | |
| **21** | Pyruvate | 2.37 | 34.2 | $CH_2$ (s) |
| **22** | Free Glycerol | 3.56 | 62.2 | $CH_2$ |
| | | 3.66 | 62.2 | $CH_2$ |
| | | 3.80 | 69.4 | CH |
| **23** | Adenine | 8.23 (s) | | |
| | | 8.35 (s) | | |

| | | | | |
|---|---|---|---|---|
| **24** | Succinate | 2.41 | 34.2 | $CH_2$ |
| **28** | Serine | 3.87 | 61 | $\alpha$-CH |
| | | 3.94 | 57 | $\beta$-$CH_2$ |
| **29** | N-acetyl | 2.06 (s) | 24.8 | $CH_3$ |
| **30** | Phosphorylcholine | 3.22 | 54.7 | $N(CH_3)_3$ |
| | | 3.61 | 67.3 | $NCH_2$ |

**Suppl. Table 2.** Genes retrieved in the "Alanine, aspartate and glutamate metabolism" and "Glutathione metabolism" KEGG pathways from the joint pathway analysis (Fig.1D) of the metabolites significantly different between CTR and NF-YA$^{KD}$ cells and datasets of differentially expressed genes after our shRNA-mediated knock-down of NF-YA in HCT116 cells (GSE70543) or siRNA-mediated NF-YA knockdown in HCT116 cells (GSE56788). Genes are listed in alphabetical order.

ALDH4A1
ALDH5A1
ASNS
CHAC1
CPS1
GLS
GLUL
GOT1
GPT2
GSS
GSTK1
GSTM1
GSTO2
GSTT2
GSTT2B
IDH2
MGST2
OPLAH
RIMKLA
RRM2
SRM

**Suppl. Table 3.** List of oligonucleotides used in RT-qPCR and ChIP-qPCR.

| Gene | Sequence (5'-3') | Application |
|---|---|---|
| NF-YA | F: GGGCTTCTGTCACCGAAAAG | RT-qPCR |
| | R: GCCGAGACTCATGCAGGTAT | |
| NF-YAl | F: CAGACCCTCCAGGTAGTCCA | RT-qPCR |
| | R: CAAACCCTGTGTTCCAGAAA | |
| NF-YAs | F: CAGAGTGGACAGGAATCTCAC | RT-qPCR |
| | R: CCTTGGACCTGCTGCTGAA | |
| ASNS | F: GCACGCCCTCTATGACAATG | RT-qPCR |
| | R: TTCAACAGAGTGGCAGCAAC | |
| GLS1 | F: CCAGAAGGCACAGACATGGT | RT-qPCR |
| | R: ACCACCATTAGCCAGTGTCG | |
| GLUD1 | F: TTGGAAAGCATGGTGGAACTA | RT-qPCR |
| | R: GCAGAACGCTCCATTGTGTAT | |
| GLUL | F: CAATGCCCGACGTCTAACTG | RT-qPCR |
| | R: GGGCTTCTGTCACCGAAAAG | |
| SLC1A5 | F: GGTTCTGGTCTCCTGGATCA | RT-qPCR |
| | R: AAGGCGGGCAAAGAGTAAAC | |
| SLC7A5 | F: TACTTCACCACCCTGTCCAC | RT-qPCR |
| | R: TGGAGGATGTGAACAGGGAC | |
| SLC38A1 | F: CAAGTCTTTGGCACCACAGG | RT-qPCR |
| | R: ACTATCACCACCAGAACGCG | |
| SLC38A2 | F: CTACTCCTACCCCACCAAGC | RT-qPCR |
| | R: GGATTCCACTGCCCACAATC | |
| CD86 | F: CTGCTCATCTATACACGGTTACC | RT-qPCR |
| | R: GGAAACGTCGTACAGTTCTGTG | |
| CD206 | F: GGGTTGCTATCACTCTCTATGC | RT-qPCR |
| | R: TTTCTTGTCTGTTGCCGTAGTT | |
| RPS20 | F: GCGCCTCTTATCAAGTCAGC | RT-qPCR |
| | R: CGGAAAAACACCCGTGGAG | |

| | | |
|---|---|---|
| GLS1 | F: CAGTTTGACTCCTCTCCCCG | qChIP |
| | R: TGGCTCAAATCCTCGACCTC | |
| GLUL | F: TTCTTTCCCAGTTCGCTTGC | qChIP |
| | R: TTAGGAGAGGAGAGGAGGCC | |

### c) Suppl. MATERIALS AND METHODS

Extensive description of methods and reagents.

#### Cell lines and lentiviral transduction

Colon cancer HCT116 cells (ATCC Cat# CCL-247) were grown in IMDM Medium (Biowest, France), colon cancer SW-480 cells (ATCC Cat# CCL-228) were grown in DMEM High Glucose Medium (Biowest, France), and human monocytic THP-1 cells (ATCC Cat# TIB-202) were grown in Roswell Park Memorial Institute culture medium (RPMI 1640, Biowest). All media were supplemented with 2 mM glutamine, 100 IU/ml penicillin, 100 µg/ml streptomycin and 10% heat-inactivated FBS (Gibco). Stable NF-YA-overexpressing cell lines were obtained by lentiviral infection of HCT116 cells as previously reported [17]. NF-YA inactivation was obtained by lentiviral infection of HCT116 cells with pLKO.1 shRNA lentiviral particles (MOI = 4) and harvested 48 h post-infection, as previously described [45]. All cells were grown at 37°C in a humidified incubator containing 5% $CO_2$ and checked for mycoplasma contamination using Mycoplasma Test Kits (Biological Industries, #20-700-20 or Lonza, #LT07-318).

#### Glutamine starvation and cell treatments

Gln depletion experiments were performed by plating cells in complete medium for 24 h, followed by washing with PBS and transfer into Gln-free medium, supplemented with 10% Dialyzed FBS (HyClone™ dialyzed Fetal Bovine Serum, #SH30079). When indicated, we concomitantly added Gln pathways inhibitors: 10 µg/ml Glufosinate-ammonium (Sigma-Aldrich, #45520), 1µM V-9302

(MedChemExpress, #HY-112683A) or 25nM Telaglenastat -CB-839- (MedChemExpress, #HY-12248).

**Cell proliferation assay**

3x10$^3$ HCT116 or SW480 cells were seeded into 96-well plate and grown in complete medium for 24h. After Glutamine starvation or cell treatment, as specified in the relevant figures, PrestoBlue reagent (#A13261, Thermo Fisher Scientific, MA) was added to the medium (1:9 v/v) and incubated for 1 h at 37°C. Cell viability was calculated by quantifying PrestoBlue reduction, measuring the absorbance at 570-620 nm, according to the manufacturer's protocol, using a GloMax Discover microplate reader (Promega). At least three independent experiments were performed.

**2D and 3D cell migration assays**

For 2D wound healing assays, HCT116 cells (1.2x10$^5$) were cultured into *Ibidi* culture-inserts (#80209, Ibidi GmbH, Germany) until confluence. After removal of the insert to create a gap, cells were gently washed with 1X PBS and Gln-free medium with or without 10 µg/ml Glufosinate was provided. Images were acquired with an EVOS M5000 microscope (Thermo Fisher Scientific, MA) immediately after insert removal (T0), after 24h (T1) and 48h (T2). Wound areas were measured (Photoshop software) and wound closure (%) was calculated with the formula 100- [(final gap area/initial gap area)×100].

For 3D spheroid-based micrometastasis assay, multicellular tumor spheroids (MTSs) were generated from 2x10$^3$ HCT116 cells by hanging-drop method, in 40 µl droplets. 72h after seeding, spheroids were collected, washed 3 times in PBS, and transferred into 48-well plates (2 MTSs/well). MTSs were then grown in complete IMDM medium or medium w/o Gln ± 10 µg/ml Glufosinate. After 10 days, MTSs and MTS-derived colonies were fixed and stained with 0.1% crystal violet solution. Plates were imaged by EVOS M5000 imaging system (Thermo Fisher Scientific, MA).

**Protein extraction and Immunoblotting**

Cells were lysed into 1X SDS sample buffer (25 mM Tris–HCl pH 6.8, 1.5 mM EDTA, 20% glycerol, 2% SDS, 5% β-mercaptoethanol, 0.0025% Bromophenol blue) and western blots were performed with nitrocellulose membrane in a Trans-Blot Turbo Transfer System (Bio-Rad, USA). The following primary antibodies (1:1000 in 1X TBS, 1mg/ml BSA) and peroxidase conjugated secondary antibodies (1:20000 in 1X TBS, 1% western blot milk) were used: anti-NF-YA (#PA528990, Invitrogen), anti-Glutamine Synthetase -GLS- (#66323-1-Ig, Proteintech), anti-Tubulin (#66031, Proteintech), anti-Glutaminase -GS- (#12855-1-AP, Proteintech), anti-Vinculin (V4504, Merck KGaA), secondary anti-mouse (#A90-116P, Bethyl Lab), and secondary anti-rabbit (#A16023, Invitrogen). Chemiluminescence signals were acquired with an Amersham Imager AI680 RGB (GE Healthcare), using Westar Supernova HRP substrates (Cyanagen) and SignalBright Max Chemiluminescent Substrate (Proteintech).

**RNA extraction and RT-qPCR**

RNA was extracted by using Riprospin II mini Kit (#314–150, GeneAll), according to the manufacturer's protocol. 200 ng of RNA was retrotranscribed with PrimeScript RT Reagent Kit (#RR037A, Takara Bio). Quantitative RealTime PCRs were performed with SsoAdvanced Universal SYBR Green Supermix (#1725274, Bio-Rad, USA) using Biorad CFX connect Real-Time PCR Detection System. Oligonucleotides sequences are listed in Suppl. Table 3. Data were analyzed using the Bio-Rad CFX Maestro 2.0 software (Bio-Rad). mRNA expression levels were normalized to Rps20 and relative fold change of transcripts were calculated with the formula $2^{-(\Delta\Delta Ct)}$, with control cell line or control growth medium used as reference sample, as indicated in figure legends.

**Promoter analysis**

In silico analysis of promoter sequences (−950 to +50 relative to transcription start site) for the presence of putative NF–Y motifs was performed with the computational algorithm Lasagna-Search 2.0 [24] (https://biogrid-lasagna.engr.uconn.edu/lasagna_search/). We performed the analysis with TRANSFAC and JASPAR core matrices for NF–Y and NFYA (TRANSFAC Accession Numbers M00185, M00209, M00287 and JASPAR ID MA0060.1) using default system parameters and cutoff p-value of 0.001. We assessed the presence of the CCAAT/ATTGG pentanucleotide by manual inspection of each promoter in the UCSC Genome Browser through the "Short Match" Track setting, and the presence of NF–Y binding from the ENCODE Project through the "Transcription Factor ChIPseq Peaks from ENCODE 3” track in the UCSC Genome Browser (human GRCh38/hg38, http://genome.ucsc.edu).

**Chromatin immunoprecipitation (ChIP)**

Chromatin was prepared from sub-confluent HCT116 cells and ChIP was performed as previously described [46] The following antibodies were added to each IP and incubated overnight at 4 °C on a rotating wheel: 1 µl of anti-NF-YA (#C15310261, Diagenode), 2 µg of anti-H3K4me3 (#C15410003, Diagenode) or control IgG (#30000-0-AP, Proteintech). Immunoprecipitated DNAs were isolated by phenol-chloroform extraction and resuspended in TE buffer. Quantitative Real-Time PCRs were performed using SsoAdvanced Universal SYBR Green Supermix (#1725274, Bio-Rad) using Biorad CFX connect Real-Time PCR Detection System. Results are presented as % of INPUT DNA, which was calculated with the formula $2^{-(\Delta Ct)}$. Sequences of primers are listed in Suppl. Table 4.

**Zebrafish Xenograft Injection of cancer cells**

Adult zebrafish and embryos of the Tg(fli1a:EGFP) transgenic strain were maintained and handled in accordance with the European Directive 2010/63/EU and Italian law (D.Lgs. 26/2014). Fertilized eggs were kept in embryo water supplemented with 0.1% Methylene Blue at 28 °C. Before manipulation, embryos were anesthetized with 0.02% tricaine (Ethyl 3-aminobenzoate methanesulfonate salt, Sigma-Aldrich® Merck KGaA).

For inhibition of Gln uptake and synthesis, NF-YAl$^{high}$ HCT116 cells were pre-treated for 2h in Gln-free medium with 10 µg/ml Glufosinate-ammonium and 1µM V-9302. Control or pre-treated cell were labelled with a red fluorescent viable dye (CellTracker Orange CMRA, Invitrogen, Carlsbad, CA, USA) and resuspended in PBS (CTR) or in 10 µg/ml Glufosinate-ammonium and 1µM V-9302 at a concentration of 2.5 × $10^5$ cells/µL. These cell suspensions were then grafted into the perivitelline space, near the sub-intestinal vein (SIV) plexus, of 48 hours post-fertilization Tg(fli1a:EGFP) embryos using a pulled micropipette. Three hours after implantation, embryos were screened to select correctly grafted larvae and to discard those displaying circulating cells. Injected embryos were maintained at 32 °C until image acquisition.

Circulating cells were monitored in vivo after 24h using a fluorescence stereomicroscope (Nikon SMZ25 equipped with NIS-Elements software), and images were subsequently analyzed with NIS-Elements (Nikon Corporation, Tokyo, Japan).

***ex vivo*** **High Resolution Magic Angle Spinning (HR-MAS) NMR**

HCT116 were harvested 48h-post infection with shScramble (CTR) or shNF-YA (NF-YA$^{KD}$) lentiviral vectors, as described above. The *ex vivo* NMR analysis was performed using 4-6 x $10^6$ HCT116 cells. One- and two- dimensional spectra were acquired. Cells were introduced in a 50 µl MAS zirconia rotor (4 mm OD) with 10 µl of deuterated water (D2O), closed with a cylindrical insert to increase sample homogeneity, then transferred into the probe cooled to 5 °C. $^1$H and $^{13}$C HR-MAS NMR spectra were recorded with a Bruker Avance III HD 600 MHz spectrometer, operating at 600.13 and 100.61 MHz, respectively. The whole experiments were performed at 5 °C to prevent cell degradation processes [47]. Samples were spun at 4000 Hz. After the set up, three different types of one and two-dimensional (1D, 2D) spectra were acquired.

The 1D 1H experiments are acquired by using: i) a composite pulse sequence (zgcppr), with 2.5 s water-presaturation during relaxation delay, 8 kHz spectral width, 32 k data points, 64 scans; ii) a watersuppressed spin-echo Carr-Purcell-Meiboom-Gill (CPMG) sequence (cpmgpr) with 1.5 s water presaturation during relaxation delay, 1 ms echo time (τ), and 360 ms total spin-spin relaxation delay (2nτ), 8 kHz spectral width, 32 k data points, 128 scans; and iii) a sequence for diffusion measurements based on stimulated echo and bipolar-gradient pulses (ledbpgp2s1d) with big delta 200 ms, eddy current delay Te 5 ms, little delta 2*2 ms, sine-shaped gradient with 32 G/cm followed by a 200 µs delay for gradient recovery, 8 kHz spectral width, 8 k data points, 256 scans. Two-dimensional (2D) COSY spectra were acquired using a standard pulse sequence (cosygpprqf) and 0.5 s water presaturation during relaxation delay, 8 kHz spectral width, 4 k data points, 32 scans per increment, 256 increments. 2D 1H,1H-TOtal Correlation SpectroscopY (TOCSY) spectra were acquired using a standard pulse sequence (mlevphpr) and 0.5 s water-presaturation during relaxation delay, 100 ms mixing (spin-lock) time, 4 kHz spectral width, 4 k data points, 32 scans per increment,128 increments. 2D 1H,13C-heteronuclear single quantum coherence (HSQC) spectra were acquired using an echo-antiecho phase-sensitive standard pulse sequence (hsqcetgp) and 0.5 s relaxation delay, 1.725 ms evolution time, 4 kHz spectral width in f2, 4 k data points, 128 scans per increment, 17 kHz spectral width in f1, 256 increments.

Areas of selected peaks of metabolites were estimated in 1D 1H CPMG spectra using Mnova software (MestReNova, ver. 8.1.0, 2012 Mestrelab Research S. L., Santiago de Compostela, Spain), with an automated fitting routine based on the Levenberg-Marquardt algorithm applied after manual peak selection, adjusting peak positions, intensities, line widths and Lorentzian/Gaussian ratios, until the residual spectrum was minimized [48]. Here we defined relative concentration as the ratio of the relative resonance area of all the quantifiable metabolites and the respective cells number. Data were reported as means ± standard errors (arbitrary units). Statistical analyses were performed using MetaboAnalyst 2.0. For Student's t-test, paired two-sample test was used to determine the means. $P < 0.05$ was considered to indicate a statistically significant difference. Suppl. Fig. 1 reports the $^1H$ spectrum with water presaturation and the cpmg spectrum. The visible and clear resonances are assigned. The major metabolites are labelled: creatine (Cr), lactate (Lac), taurine (Tau), macromolecules (MM), acetate (Ac), glutamate (Glu), glycine (Gly), (BCAA).

Metabolite set enrichment analysis and integrated joint pathway analysis were performed using MetaboAnalyst 6.0 [49] (https://dev.metaboanalyst.ca/MetaboAnalyst/), using KEGG pathway-based analyses.

For enrichment analysis, we used metabolite sets containing at least 5 entries. For integrated pathway analysis, we used the significantly different metabolites between CTR and NF-YA$^{KD}$ cells together with our matched transcriptomic dataset from shRNA-mediated NF-YA$^{KD}$ HCT116 cells (GSE70543) or with an independent public dataset of siRNA-mediated downregulation of NF-YA in HCT116 cells (GSE56788). Integrated pathway analysis was performed with Betweenness Centrality topology analysis (measure of the number of shortest paths from all nodes to all the others that pass through a given node) and combined p values integration method (enrichment analysis is performed separately for genes and metabolites in their "individual universe", and then individual p-values are combined via weighted Z-tests).

#### Metabolic analysis using seahorse technology

Metabolic characterization of HCT116 cells was performed using a Seahorse XFe24 Extracellular Flux Analyzer (Seahorse Bioscience). Briefly, 2.5 × $10^4$ cells were seeded into the Seahorse XF Cell Culture 24well-plate (Agilent Technology) one day before the experiment. The raw data from the Seahorse analysis were normalized to the number of seeded cells, in order to minimize variation from well-to-well due to potential non-uniform cell seeding across the microplate. After Seahorse metabolism analysis, the cells were stained with Hoechst3342 and then imaged using an ImageXpress MicroConfocal fluorescence microscope (Molecular devices). The number of nuclei was then counted using the Metaxpress Software (Molecular Devices), allowing an automatic and precise direct cell count. The Mito Fuel Flex Test was performed according to the manufacturer's instructions to measure the OCR and test dependency, capacity and flexibility of cells to oxidize glucose, glutamine or long-chain fatty acids as mitochondrial fuels. Metabolic parameters were exported and calculated according to the manufacturer's instructions (Agilent Technologies) using the Seahorse Wave software (Agilent Technologies).

#### Detection of mitochondrial superoxide (MitoSOX)

5 × $10^3$ HCT116 cells/well were seeded in 96-well plate for 24h, then incubated in Gln-deprived medium with or without 50μM H2O2 for 72h. Mitochondrial superoxide level was measured with the MitoSox red dye (Thermo Fisher Scientific, Inc.) according to the manufacturer's procedures. Fluorescence was measured after 30 min using a GloMax Discover microplate reader (Promega).

#### Measurement of glutamine and GSH

HCT116 cells were seeded at 1 x $10^4$ cells/well in a 96-well plate for 24h, then treated with complete or Gln-depleted medium for 24h. Gln was measured in cell lysates and cell growth media using the bioluminescent Glutamine/Glutamate-Glo Assay kit (Promega, Cat#: J8021), while total Glutathione was quantified into cells with a Colorimetric Assay Kit (elabscience, Cat#: E-BC-K097-M), according to manufacturers' instructions. Absorbance and luminescence were measured using a GloMax Discover microplate reader (Promega).

**Microfluidic circulatory system fabrication and circulation of HCT116 colon cancer cells**

We developed a circulating system exploiting a peristaltic pump for which a minimum flow rate of 0.33 ml/s was possible. The system, apart from the regions inside the peristaltic pump and the region for cell injection (see Fig. 5C), exploits a tube with an internal diameter of 0.75 mm. These values allow us to evaluate the condition of laminarity of the flow in our system. Indeed, calculating the Reynolds number $Re = \frac{\rho v D}{\eta}$, where $\rho$ is the density of the culturing medium, which can be approximated to 1 g/cm$^3$, $v$ is the velocity of the flow, $D$ is the diameter of the tube (the main tube) and $\eta$ is the viscosity of the culture medium, which can be approximated as $\eta = 10^{-3}$ Ns/m$^2$, we obtain $Re = 562$, which is below the threshold of 2200 for turbulent flow. Shear stress is maximum near the surface of the tube and minimum in the central region changing linearly as a function of the distance from the central region of the tube, so, we calculated the maximum value according to the Poiseuille law:

$$\tau_{max} = \frac{4Q\eta}{\pi R^3}$$

where $Q$ is the flow rate and $R$ the radius of the main tube. For the maximum shear stress value, we obtain: $\tau_{max} = 31.3$ dyne/cm2, a typical value for arterial flow or capillary bed. For cells in suspension, the relevant physical parameter is the shear rate related to a velocity gradient. This quantity changes linearly with the distance $r$ from the central tube axis according to the formula $\tau_{max} = \tau_{max}(r/R)$ where $R$ is the radius of the tube. Accordingly, the minimum value of shear stress can be calculated using the previous formula where $r$ is the average radius of cells in suspension and we obtain a value of about 0.58 dyne/cm$^2$.

All tubing and microfluidic devices were sterilized with 70% ethanol and washed with PBS before each experiment, and air bubbles in the microfluidic system were removed during the washing steps. HCT116 cells at ~75% confluence were pre-treated for 1h in Gln-free IMDM, collected by trypsinization and suspended in Gln-free IMDM, with or without the addition of 10 µg/ml Glufosinate. 1.5 mL of HCT116 cell suspension at a concentration of 5 × 10$^5$ cells/ml were injected into the microfluidic circulatory system and recovered after 1h of circulation in Gln-free medium. Cell number was determined using a NucleoCounter NC-100 instrument (ChemoMetec).

For experiments of cell-recovery, 500µl of post-circulation cell suspension were plated in Gln-free IMDM for 24h in standard cell culture dishes, then cell number was determined. To account for any changes in viability due only to Gln-depletion, the initial pre-circulation HCT116 cell suspension was used as control for counting and plating in recovery experiments.

**THP-1 monocytes differentiation into macrophages**

Human THP-1 monocytes were differentiated into macrophages by incubation in conditioned-medium of HCT116 cells. Conditioned medium (CM) was obtained by seeding 1.5 x $10^5$ HCT116 cells/well in 6-wells culture plates: after 18h, the medium was replaced with new complete or Gln-deprived IMDM for 24h. This CM was collected, filtered through 0.45 µm syringe filter and used for THP-1 activation. THP-1 monocytes were incubated for 48h in a 50% mixture of filtered conditioned medium (cm) from HCT116 cells and RPMI with either complete or Gln-deprived supplements. Where indicated, 10 µg/ml Glufosinate and 1µM V-9302 were added in the Gln-depleted activation medium for treatment of THP-1.

Activated monocytes became adherent and the amount/viability of cells and the expression of macrophage markers by RT-qPCR were analyzed as described above.
